# Modeling the rhythmic complexity of professional drumming with an oscillation-driven reservoir computer

**DOI:** 10.1101/2024.07.17.603863

**Authors:** Yuji Kawai, Shinya Fujii, Minoru Asada

## Abstract

Musical performances, particularly in drumming, are characterized not only by their structured rhythmic patterns but also by the subtle variations in timing and amplitude series that create expressive complexity. This study proposes a neural-inspired computational model to investigate how the brain might learn and internalizes such complex rhythms. Inspired by the established roles of the cerebellum and basal ganglia in production of rhythms and timings, we utilize an oscillation-driven reservoir computer, a recurrent neural network model for temporal learning, to simulate the generation of human-like expressive drumming performances. First, the model was trained to replicate Jeff Porcaro’s distinctive hi-hat patterns. Analyses revealed that the outputs of the model incorporating high-frequency oscillators ([50, 100] Hz), closely matched the original drumming, reproducing its characteristic fluctuations and patterns in inter-beat timings (microtiming) and amplitudes. Next, the model was trained to generate multidimensional drum kit performances for various genres (funk, jazz, samba, and rock). The model’s outputs exhibited timing deviation and audio features characteristic of the original performances. Our findings demonstrate that oscillation-driven reservoir computing can replicate the rhythmic complexity of professional drumming, suggesting it as a potential computational principle for motor timing and rhythm generation. This approach provides a powerful framework for understanding how the brain generates and processes intricate rhythmic patterns.

## Introduction

While a substantial body of research has explored the mechanisms of rhythm perception [1–4], how humans learn and generate complex rhythms remains poorly understood. This generative ability allows us to form internal representations of rhythmic structures, enabling the creation of novel yet stylistically coherent musical rhythms that go beyond simple imitation [5]. Professional drumming exemplifies this complexity, incorporating subtle timing fluctuations and rich acoustic features.

The timing and velocity of beats in the performances of professional musicians include fluctuations [6–9], which create pleasurable emotional and behavioral responses [10, 11]. Such a slight timing deviation is termed as “microtiming,” which usually less than 50 ms [12, 13]. Räsänen et al. [8] extracted Jeff Porcaro’s hi-hat playing sound from Michael McDonald’s “I Keep Forgettin’ “ [14] and analyzed the hi-hat amplitudes and inter-beat intervals, i.e., time intervals between percussion strokes or hits. The timing intervals drift from the 16th note pattern and follow a Gaussian-type distribution, indicating expressive microtiming. Furthermore, detrended fluctuation analysis (DFA) [15] revealed that, like 1*/f* fluctuations, the fluctuations of the timing intervals and amplitudes have persistent correlations over long time scales, i.e., data points separated by long time intervals are not independent but exhibit correlated dependence. However, it is debated whether such microtiming induces a sense of groove (a pleasurable desire to move in response to music [16–19]) in listeners [20].

In addition to microtiming, other audio features related to a groove sensation have also been investigated. Madison et al. [17] found that beat salience (a measure of rhythmic periodicity) and event density (a measure of the variability in the event onset velocity signal) correlate with groove ratings. Stupacher et al. [21] corroborated Madison et al. [17]’s result and reported that audio signal intensity, its variability, and attack slope of onsets increase groove ratings.

There is considerable evidence that auditory-motor brain regions activate when listening to music and rhythm [1, 20, 22–27]. Kasdan et al. [27] conducted a meta-analysis of 30 functional magnetic resonance imaging studies that investigated musical rhythm processing. They revealed a large network involving the auditory and motor regions, including the bilateral superior temporal cortices, supplementary motor area, basal ganglia, and cerebellum. In particular, cerebellar activity is greater when the rhythm is more complex. It has been suggested that there is a close relationship between the perception and production of rhythms and that there is overlapped neural circuits [25, 28]. The subcortical areas, basal ganglia, and cerebellum respond to auditory time perception and production [24, 25, 29], and patients with focal lesions in these areas show reduced tracking at the beat frequency [30]. Furthermore, it has been reported that musical training induces plastic changes in the cortical auditory-motor regions, as well as in the cerebellum and basal ganglia [31]. From a broader perspective, these areas play important roles in motor learning and control [32–35]. The basal ganglia perform reward-based learning with dopaminergic modifications while the cerebellum learns by adjusting its connections (long-term depression: LTD) based on feedback (error signals) from climbing fibers. These signals help fine-tune motor skills and coordination [33]. Through these learning mechanisms, these regions can also learn motor timing [36–38]. Although the functional segregation of the cerebellum and basal ganglia in timing perception has been debated [24, 39–41], we believe that these regions share computational principles that underlie the learning of timing and rhythms.

Reservoir computing, a type of recurrent neural network (RNN), was used to model the cerebellum [42–44] and basal ganglia (striatum) [45–47]. Standard reservoir computing consists of a fixed randomly connected RNN, termed a reservoir [48, 49]. The input to the reservoir network induces complex neural dynamics, and its network states are integrated using a weighted sum (readout). The readout weights are modified to obtain the desired output, for which the least-squares method is often used. In cerebellar models, granule cells perceive cortical inputs and form an RNN (reservoir) with the Golgi cells, and readout training is realized through Purkinje cell plasticity (LTD). In the basal ganglia models, the striatum receives cortical inputs and dopamine modification in the striatum is regarded as readout training. Reservoir computing, which enables temporal information processing based on neural dynamics, has been utilized as a model to explain motor timing learning [47, 50–54]. In the motor timing task, there was a stimulus-free interval between stimuli. Therefore, the neural activity in the reservoir elicited by the initial onset signal must be sustained for an interval without any input. Vincent-Lamarre et al. [51] demonstrated that providing a reservoir with sinusoidal oscillations allows the reservoir to sustain their activity during an interval, allowing timing to be successfully learned. Kawai et al. [54, 55] improved performance by adding output feedback to the model and showed that it could learn and predict complex time series. To date, however, reservoir models have not learned and generated real-world rhythms.

This study investigates the computational principles underlying the learning and generation of complex drum rhythms. To this end, we employ the reservoir model proposed by Kawai et al. [54], a framework known as a timing-learning machine. The readout was trained with the desired target time series from the encoded performances of professional drummers. By feeding back the output of a reservoir to its input, the reservoir learns the next output from the current output; i.e., it predicts one time ahead. When generating rhythms, the reservoir repeats this one-time-ahead prediction and provides feedback to generate a time series. In standard reservoir models, the generated time series tends to be simple and periodic. To represent fluctuating timing intervals and amplitudes of skilled drumming performances, we input oscillations with multiple frequencies to the reservoir. It is expected that the input of oscillation superimposition produces more complex dynamics than the network-inherent dynamics, which facilitates the learning of acyclic drumming. This model suggests that oscillation-driven reservoir computing is a common computational principle used for timing and rhythm generation. Model outputs are then analyzed to estimate to what extent they mimic the original performances with respect to microtiming and audio features that related to a groove sensation.

First, the model learns the one-dimensional hi-hat rhythm pattern by Jeff Porcaro [8]. We tested the models with oscillations in different frequency bands and analyzed the nature of the generated time series. Using the same analyses as Räsänen et al. [8], we examined whether the rhythms generated after learning were similar to the original performance. Next, the model was trained to perform as a multidimensional drum kit. Microtiming and audio features of the outputs were analyzed and compared with those of the original performances. We present examples of the generated time series to demonstrate that the reservoir model can copy and generalize the performances in different genres.

### Related studies

Entrainment [56–58] and error-correction [59–62] models have been proposed to explain the ability to perceive a beat from a computational perspective (for a review, see Ref [4]). The entrainment models rely on oscillators that resonate and entrain their inputs. Entrainment refers to the synchronization of two or more oscillating systems as a result of an external forces or interactions between them. Beat perception is regarded as synchronizing an external rhythmic input with a preprepared oscillators. Classical error-correction models adjust the timing of the next finger tap to reduce the error between the current tap and the stimulus [59]. Bose et al. [60] proposed an error-correction model in which a beat-generation oscillator learned the spiking phase and period of a stimulus neuron receiving an isochronous stimulus sequence. A comparator counts the number of spikes in the oscillator and stimulus neuron and adjusts the oscillator frequency according to the error between their counts. Egger et al. [61] derived an error-correction model using a firing-rate neuron model. The neurons build an oscillator that activates a ramp function, and its value is reset when a certain threshold is exceeded. The ramping speed depends on an input parameter that controls the oscillator frequency. The input value was adjusted at each reset such that the oscillator frequency matched the time between successive stimuli. These models represent isochronous rhythms by tuning the frequency of an oscillator or the frequencies of oscillators. Simple oscillators can only represent simple beats and do not generate complex rhythms. The reservoir model used in this study is an error-correction model because it adjusts the readout based on errors. However, the reservoir network transforms the oscillation inputs into complex dynamics with large dimensions, which generates complex rhythms.

Machine learning methods for generating music and complex rhythms are being actively developed [63, 64]. From reviewing the generation of drum performances, Hutchings [65] proposed a method for generating a full drum kit part for a provided kick-drum sequence using an RNN sequence-to-sequence method. Makris [66] combined a long short-term memory- (LSTM) based architecture with a feedforward neural network module. The former module learns the sequences of successive drum events, whereas the latter deals with musical information on the guitar, bass, beat structure, and tempo. Thus, it can generate rhythms conditioned by musical information. Variational autoencoders (VAEs) learn to compress the input data into a latent distribution and reconstruct their input data [67]. The decoder network generates a novel output from the latent space. GrooVAE [68] automatically composes drum performances using a VAE, which generates and controls expressive drum performances. Furthermore, Transformers [69], which are a type of a deep learning model, have been applied in the field of music generations (e.g., [70–72]). The learning algorithms of these machine learning systems [63–72] are based on error back-propagation and do not aim to understand the brain processing. Their learning costs are very high, and they require large computational resources and training data. In contrast, reservoir computing is more biologically plausible because its learning algorithm does not rely on backpropagation and can be realized by simple rules, including LTD and dopaminergic modification in readout. Reservoir computing has fewer training parameters and can, therefore, learn from a small amount of data. This study is the first to use reservoir computing to generate drum performances.

## Methods

### Oscillation-driven reservoir computing [54]

The reservoir is a randomly connected RNN with firing-rate neural units that receive multiple sinusoidal oscillator inputs (Fig 1). The frequencies and phases of the oscillators are randomly determined, and once set, they are fixed across all trials. The initial states of the reservoirs are randomly determined. To reduce the initial state dependency, an onset signal is fed into the reservoir at time zero. This signal is a short, strong single pulse that resets the reservoir activity [50, 52]. The output is obtained by the linear summation (readout) of the reservoir activity. The output is then fed back into the reservoir at the next time step. The readout weights are trained to minimize the errors between the outputs and target time series using recursive least squares [73], which is an online learning method. The other connection weights for the oscillator inputs, onset signal input, reservoir, and output feedback are determined randomly and fixed through learning.

**Fig 1.**
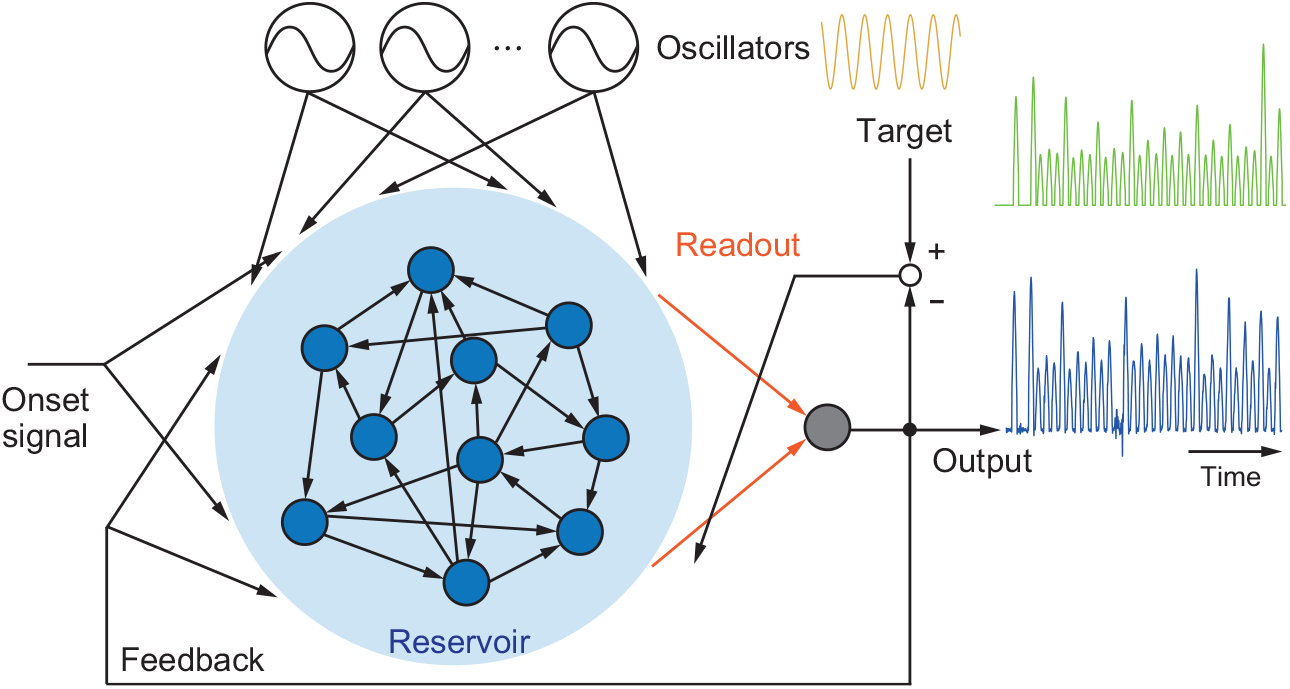
Oscillation-driven reservoir computing. Sinusoidal oscillators drive a reservoir, which is a randomly connected recurrent neural network with fixed connection weights. The output is given as a linear summation of the reservoir states (readout, shown as orange arrows), which is fed back into the reservoir. Only the readout weights are modulated using recursive least-squares. Adapted from Ref. [54].

The reservoir is a network with *N* neural units whose state vector is represented as **x**(*t*) = (*x*_1_, *x*_2_, …, *x*_*N*_)^⊤^ at time *t*. The initial values of **x**(*t*) are randomly drawn from a uniform distribution of [−1, 1]. Its network dynamics follows

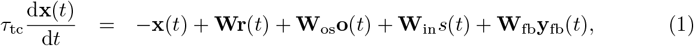

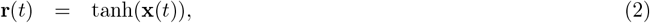

where *τ*_tc_ denotes the time constant, and **W, W**_os_, **W**_in_, and **W**_fb_ respectively denote the reservoir recurrent weight matrix, oscillator input weight matrix, onset signal input weight vector, and feedback weight matrix. The variables **o**(*t*), *s*(*t*), and **y**_fb_(*t*) respectively denote the oscillation inputs, an onset signal, and the output feedback signals.

**W** is an *N* × *N* weight matrix, each component of which has a non-zero value with a probability *p*. The non-zero values are drawn from a Gaussian distribution with a mean of zero and a standard deviation of 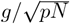, where *g* is the gain of the reservoir weights. The oscillators are *N*_os_ sinusoids. The vector of the oscillator input is represented as 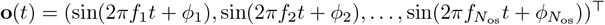, where *f*_*i*_ and *ϕ*_*i*_ respectively denote the frequency and the random initial phase. The frequencies are randomly determined from a uniform distribution of [*f*_min_, *f*_max_] Hz. Parameters *f*_*i*_ and *ϕ*_*i*_ are fixed during learning and testing. **W**_os_ is an *N*_os_ × *N* matrix for the oscillator input to the reservoir. The scalar onset signal *s*(*t*) from −50 ms to 0 ms, is 1 and at other times, 0. **W**_in_ is the weight vector of size *N* for the onset signal inputs into the reservoir. The output is fed back into all reservoir units as **y**_fb_(*t*). The feedback weights **W**_fb_ is an *N*_ro_ × *N* matrix. The components of **W**_os_, **W**_in_, and **W**_fb_ are drawn from Gaussian distributions with a mean of zero and standard deviations of 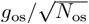, and *g*_in_, 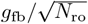, where *g*_os_, *g*_in_, and *g*_fb_ denote the gains of the oscillation input weights, onset signal input weights, and feedback weights, respectively.

To obtain the *i*th output *y*_*i*_(*t*) (*i* = 1, 2, · · ·, *N*_ro_), **r**(*t*) is weighted as

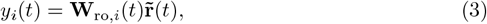

where **W**_ro,*i*_(*t*) denotes the *i*th (3*/*2 ∗ *N*) × 1 readout weight vector, and 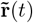 is the concatenation of **r**(*t*) and a vector of half components of **r**(*t*) squared. This has been empirically shown to improve forecasting performance [74–76]. **W**_ro,*i*_(*t*) has an initial value for the zero matrix and is modulated by recursive least-squares, which is an online learning method. At time *t*, 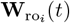 is updated as follows:

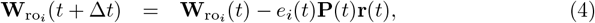

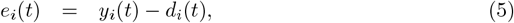

where *d*_*i*_(*t*), *e*_*i*_(*t*), and **P**(*t*) respectively denote the *i*th target, the *i*th error, and (3*/*2 ∗ *N*) × (3*/*2 ∗ *N*) matrix. **P**(*t*) is updated as

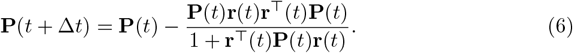

The initial value of **P**(*t*) is given as (1*/a*)**I**, where **I** denotes the identity matrix, and *a* is a constant.

In the simulations, the numerical solutions to Eq (1) were obtained using the Euler method, where the simulation step size Δ*t* was set to 1 ms. The recursive least-squares (Eqs (4)–(6)) was applied once every two steps, and Δ*t* was set to 2 ms. The simulation started at −250 ms, and the training period was from 1 ms to 20,000 ms. The training was repeated ten times, and the performance was evaluated during the untrained testing period.

We used the parameter values listed in Table 1 in all the simulations. These parameters follow those in Ref [54]. To learn the hi-hat-only performance, we set *N*_ro_ = 1; to learn the funk performance, *N*_ro_ = 3; and to learn other drum-kit performances, *N*_ro_ = 5.

**Table 1.** Parameter settings.

| Parameter | Description | Value |
| --- | --- | --- |
| $N$ | Reservoir network size | 5000 |
| $N_{\text{os}}$ | Number of oscillators | 10 |
| $g$ | Gain of reservoir weights | 1.2 |
| $g_{\text{os}}$ | Gain of oscillation input weights | 0.5 |
| $g_{\text{in}}$ | Gain of onset signal input weights | 5 |
| $g_{\text{fb}}$ | Gain of feedback weights | 3 |
| $p$ | Connection probability | 0.1 |
| $\tau$ | Time constant | 10 ms |
| $a$ | Initial value for recursive least squares | 1 |

### Detrended fluctuation analysis

DFA is a statistical method widely used to analyze the presence of long-range correlations in time-series data, particularly for non-stationary data, such as heartbeat time series [15], brain oscillations [77, 78], and human musical rhythms [8, 79]. Long-range correlations refer to the persistence of dependencies between values that are far apart in a sequence. These correlations indicate that values separated by a long time period are still statistically related, which can indicate the underlying processes or structures driving the data.

We applied DFA to the inter-beat intervals and amplitudes. Let *u*(*i*) be the *i*th hit time, and the *i*th inter-beat interval be denoted by *τ* (*i*) = *u*(*i* + 1) − *u*(*i*). Then, we calculate the fluctuation of the timing interval from the mean ⟨*τ* ⟩, i.e.,

Δ*τ* (*i*) = *τ* (*i*) − ⟨*τ* ⟩. The fluctuations are integrated as

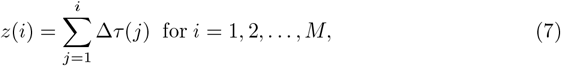

where *M* is the number of data points. Next, the integrated time series is divided into boxes of equal length *s*. Within each box, a least-squares line (*z*_*s*_(*i*)) is fitted to the data points (*y*(*i*)) to capture the linear trend. The trend represented by *z*_*s*_(*i*) was subtracted from the original data (*z*(*i*)) within the box, resulting in the detrended fluctuation: (*z*_detrended_(*i*) = *z*(*i*) − *z*_*s*_(*i*)). Finally, the root-mean-square fluctuations are calculated for each box by the following

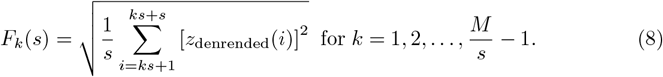

Finally, *F*_*k*_(*s*) is averaged over all *M/s* elements to obtain *F* (*s*) = ⟨*F*_*k*_(*s*) ⟩. This procedure is repeated for different box size *s*. In the presence of scaling, the relationship between *F* (*s*) and *s* may be described by *F* (*s*) ∝ *s*^*α*^, where the scaling exponent *α* is the slope of the line relating log *F* (*s*) and log *s*. Eq (7) and below applied similarly to the amplitude data series.

The scaling exponent *α* indicates the nature of the correlations; when *α* = 0.5: the series is uncorrelated, e.g., for white noise; when 0.5 *< α <* 1: the series has long-range positive correlations; when 0 *< α <* 0.5: the series has long-range negative correlations.

### Encoding and decoding of drum performances

To learn the hi-hat performance, we used a dataset of hi-hat hit timings and their amplitudes [8]. This data was obtained by applying a 100th-order FIR filter with a cutoff frequency of 8 kHz to the original sound source to extract only the hi-hat sound and then applying an onset detection algorithm [8]. To encode this data into the target time series, we first added 0.2 to all amplitudes to emphasize small amplitude hits. Single Gaussian pulses centered on the hit timing were then placed on a time series. The Gaussian amplitudes were the combined amplitudes of the hits, and their standard deviations were set to 30 ms. We cut out the first 20 s of the beginning and normalized it for the maximum and minimum to be 1 and 0 (see Fig 2).

**Fig 2.**
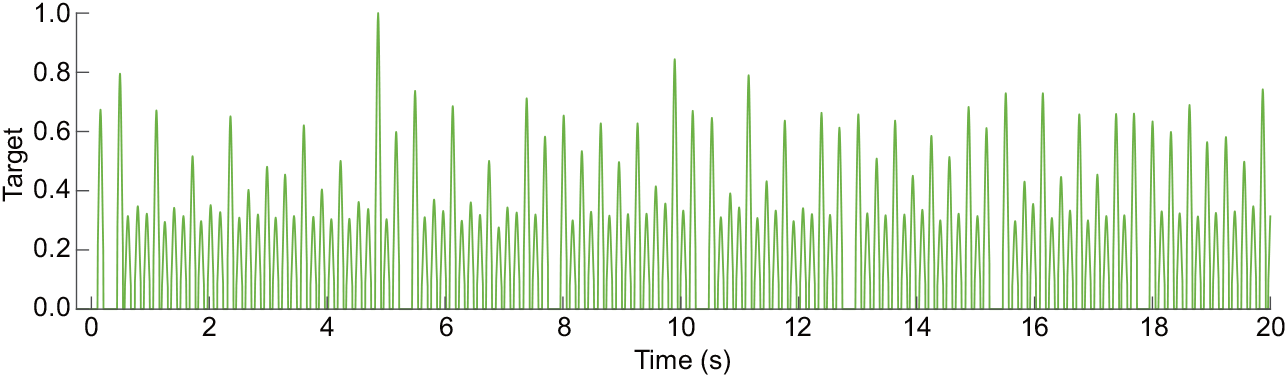
Target 20-s time series obtained from the hi-hat performance.

To convert the model output into sound data, a moving average filter was first applied to the output time series and a maximum value detection was performed to obtain hit timings and their amplitudes. The amplitudes were renormalized such that the maximum and minimum amplitudes were those of the original. Hits with amplitudes below 0.1 were ignored. We then placed hi-hat audio data at the timing points, and their intensity (loudness) was linearly modulated by the amplitudes.

We used the Groove MIDI Dataset [68] which is a large dataset comprising 1150 MIDI files of drum performances by ten professional drummers in various genres.

From this dataset, we selected four performances.

- Funk: drummer1/session1/1_funk 80 beat_4-4.mid,
- Jazz: drummer1/session1/4_jazz-funk 116_beat 4-4.mid,
- Samba: drummer1/session1/28_latin-samba_116 beat 4-4.mid,
- Rock: drummer1/session3/1_rock-prog 125_beat 4-4.mid.

Their MIDI files distinguishes between different types of drumming, e.g., between an open or closed hi-hat; however, hi-hats, snares, toms, and cymbals were grouped into one dimension each owing to the infrequency of hits. This procedure reduced the numbers of dimensions (pitches) of funk, jazz, samba, and rock respectively from seven to three, from 12 to five, from six to five, and from six to five. Similar to above, the model outputs were converted to data of peak timings and their amplitudes.

Fluctuation in the peaks resulted in small hits with small timing intervals to appear. Therefore, hits with timing intervals of 5 ms or less were removed as noise. The data were converted to the MIDI format and then converted to audio data.

In the audio analysis, we used Musical Information Retrieval (MIR) Toolbox for MATLAB [80]. The MIR Toolbox functions we used for the analysis of the six features are listed below.

- Root mean square (RMS) energy (*mirrms*): global signal intensity (RMS of the amplitude).
- Spectral flux: averaged over the entire period of a spectral flux time series as the output of *mirflux*.
- Attack (*mirpulseclarity* using ‘Attack’ option): mean attack (amplitude) slope of all hit onsets.
- Pulse clarity (*mirpulseclarity* using ‘MaxAutocor’ option): clarity of rhythmic or metrical pulsation.
- Event density (*mireventdensity*): the number of events per second.
- RMS standard deviation (SD): SD over the entire period of an RMS time series as the output of *mirrms*.

## Results

### Learning the hi-hat performance

The target time series was encoded from the hi-hat performance by Jeff Porcaro [81] in “I Keep Forgettin’ “ [14] by Michael McDonald. We used the dataset of hi-hat hit timings and their amplitudes available in Ref [8]. We encoded these data into a target time series (see the Methods section, Fig 2). These hi-hat performance data, which consist of 16th notes, have six 16th rests after 4 s, and bar breaks after these rests. The relevant audio file is attached as Audio S1.

The reservoir computing model was trained with the target 20-s time series and generated 40-s outputs. During the generation, the model was not given the target and generated outputs autonomously. The latter 20 s, in particular, are an inexperienced period during which the model is expected to produce outputs analogous to the target; that is, it generalizes the target. Fig 3A shows an example of the output generated by reservoir computing without oscillation inputs. The output fell into a completely periodic orbit and did not have the rests and fluctuations characteristic of the original performance. The audio file of the example output shown in Fig 3A is attached as Audio S2.

**Fig 3.**
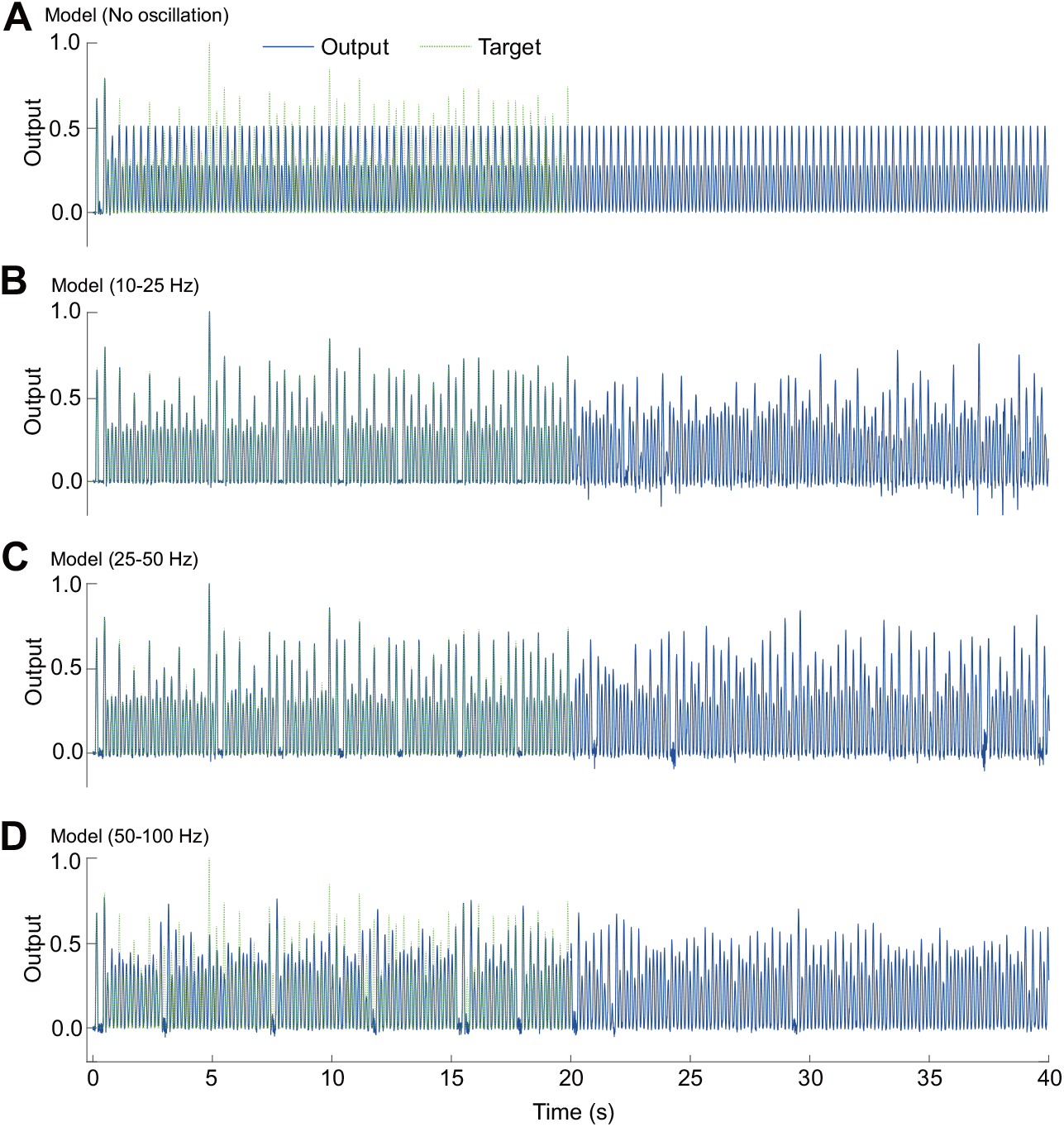
Example outputs of the reservoir model trained with the hi-hat time series. The solid blue and dashed green curves represent the model output and target respectively. A. Output of the reservoir model without oscillation inputs. B–D. Outputs of the reservoir models respectively with [10, 25] Hz, [25, 50] Hz, and [50, 100] Hz, oscillation input ranges.

In contrast to its periodic output, oscillations in the reservoir enabled it to reproduce and generalize the target (Fig 3B–D). The behavior of the outputs depended on the frequency bands of the oscillations. When oscillations of [10, 25] Hz were given to the reservoir (Fig 3B), the model reproduced the target during the first 20 s whereas the output was disturbed and lacked generalization ability in the second half. Fig 3C shows the output of the model with [25, 50] Hz oscillators. In the first half, the model followed the target; in the second half, the output was analogous to the target. When further high-frequency oscillations ([50, 100] Hz) were applied to the reservoir, the model did not replicate the target perfectly but produced an output similar to that of the target (Fig 3D). The respective audio files are attached as Audios S3 to S5.

We quantified the similarity between the model-generated outputs and the original (target) data regarding fluctuations and patterns of inter-beat timing intervals and amplitudes. In the analyses, the original data for all times of one song were used. While all the amplitude data were analyzed, only the 16th note intervals were analyzed. Therefore, only the timing interval data between 100 ms and 200 ms were analyzed. The model generated a 200-s output, the first 20 s of which were discarded, and the remaining 180-s portion was included in the analyses. The results of these analyses were averaged among the outputs of 20 models with different random seeds.

Histograms of the timing intervals of the hi-hat hits are shown in Fig 4. The timing intervals of the original data followed a Gaussian-type distribution, with a mean of 156.6 ms and a standard deviation of 8.7 ms (Fig 4A). In contrast, the output of the model without oscillators showed minimal fluctuation, with timing intervals at two discrete values (Fig 4B). Introducing low-frequency oscillations ([10, 25] Hz) induced temporal variability, but the resulting distribution was non-Gaussian and characterized by heavier tails (Fig 4C). Conversely, the models incorporating high-frequency oscillators ([25, 50] Hz and [50, 100] Hz) successfully generated interval distributions that resembled a Gaussian shape, similar to the original data (Fig 4D and E). These results demonstrate that the high-frequency model could replicate the timing fluctuations of the original performance, known as microtiming. The mean and standard deviation of the timing intervals, obtained from 20 runs for each condition, are summarized under the “*τ* (ms)” column in Table 2.

**Table 2.** Metrics of the original and model outputs. Mean (standard deviation).

| Source | $\tau$ (ms) | DFA $\alpha_\tau$ (short) | DFA $\alpha_\tau$ (long) | DFA $\alpha_A$ | $r$ |
| --- | --- | --- | --- | --- | --- |
| Original | 157 (8.7) | 0.31 | 0.72 | 0.63 | -0.48 |
| No oscillation | 155 (5.3) | 0.020 (0.025) | 0.000 (0.002) | 0.007 (0.004) | -0.75 (0.27) |
| 10–25 Hz | 151 (15.1) | 0.52 (0.033) | 0.53 (0.092) | 0.48 (0.044) | -0.075 (0.057) |
| 25–50 Hz | 156 (8.6) | 0.37 (0.068) | 0.43 (0.088) | 0.43 (0.066) | -0.34 (0.087) |
| 50–100 Hz | 155 (10.2) | 0.29 (0.068) | 0.39 (0.13) | 0.41 (0.085) | -0.48 (0.14) |

**Fig 4.**
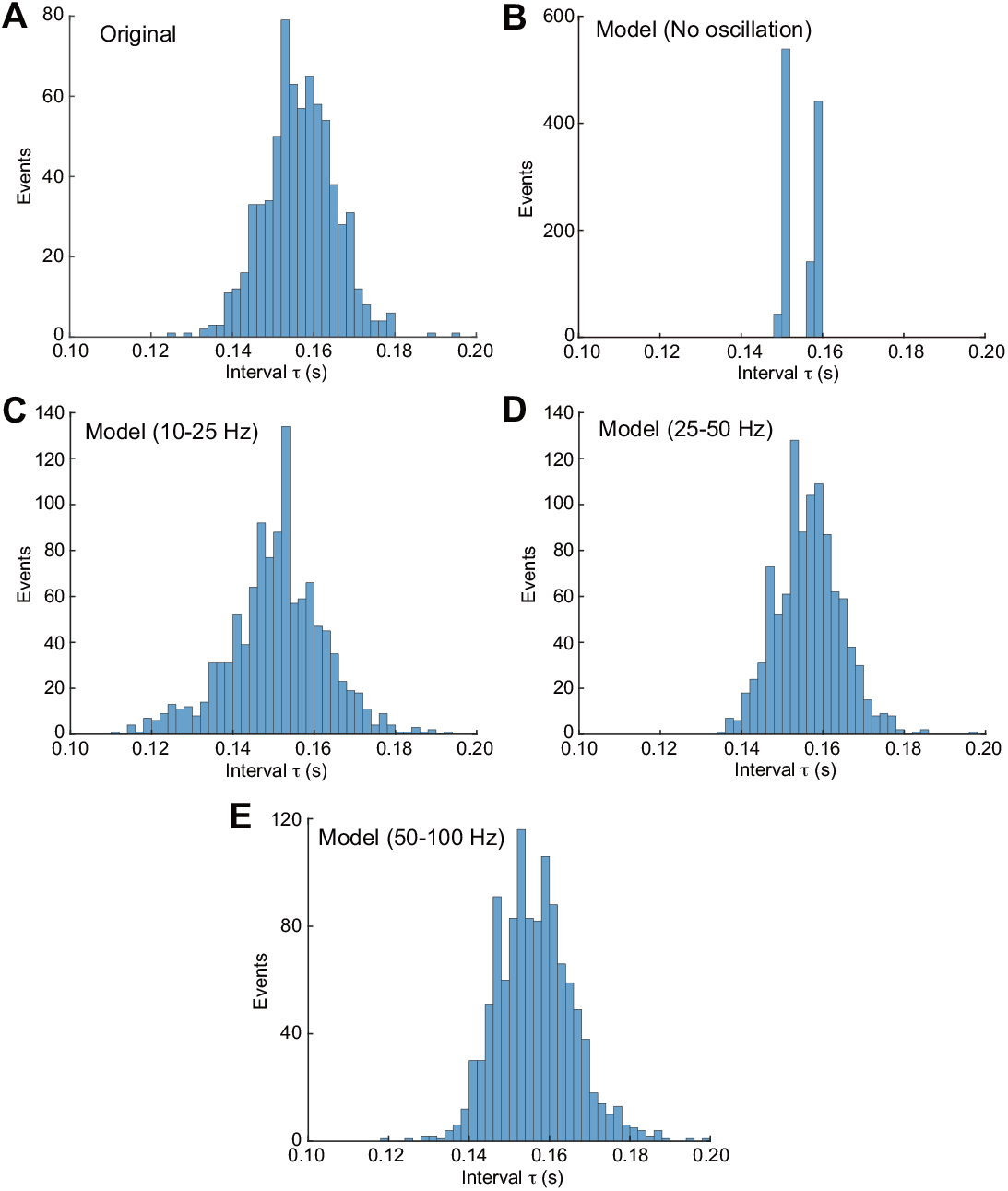
Histograms of hi-hat timing intervals. A: the original performance data. B: the model output without input oscillators. C, D, E: the model outputs with oscillators in the frequency ranges of [10, 25] Hz, [25, 50] Hz, and [50, 100] Hz, respectively.

Fig 5 shows amplitude histograms of the hi-hat hits. The original performance data displayed a bimodal distribution, reflecting a natural pattern of strong and weak accents (Fig 5A). In contrast, the model without oscillators produced amplitudes concentrated at two discrete values, resulting in two sharp peaks and a lack of variability (Fig 5B). The models with low-to mid-frequency oscillators ([10, 25] Hz and [25, 50] Hz) generated unimodal amplitude distributions that were skewed towards larger amplitudes (Fig 5C and D), failing to reproduce the accent structure. Conversely, the high-frequency model ([50, 100] Hz) produced a bimodal amplitude distribution with significant variability, closely resembling that of the original performance (Fig 5E). This result indicates that the high-frequency model successfully captured both the strong-weak accentuation and its variability as observed in the original performance.

**Fig 5.**
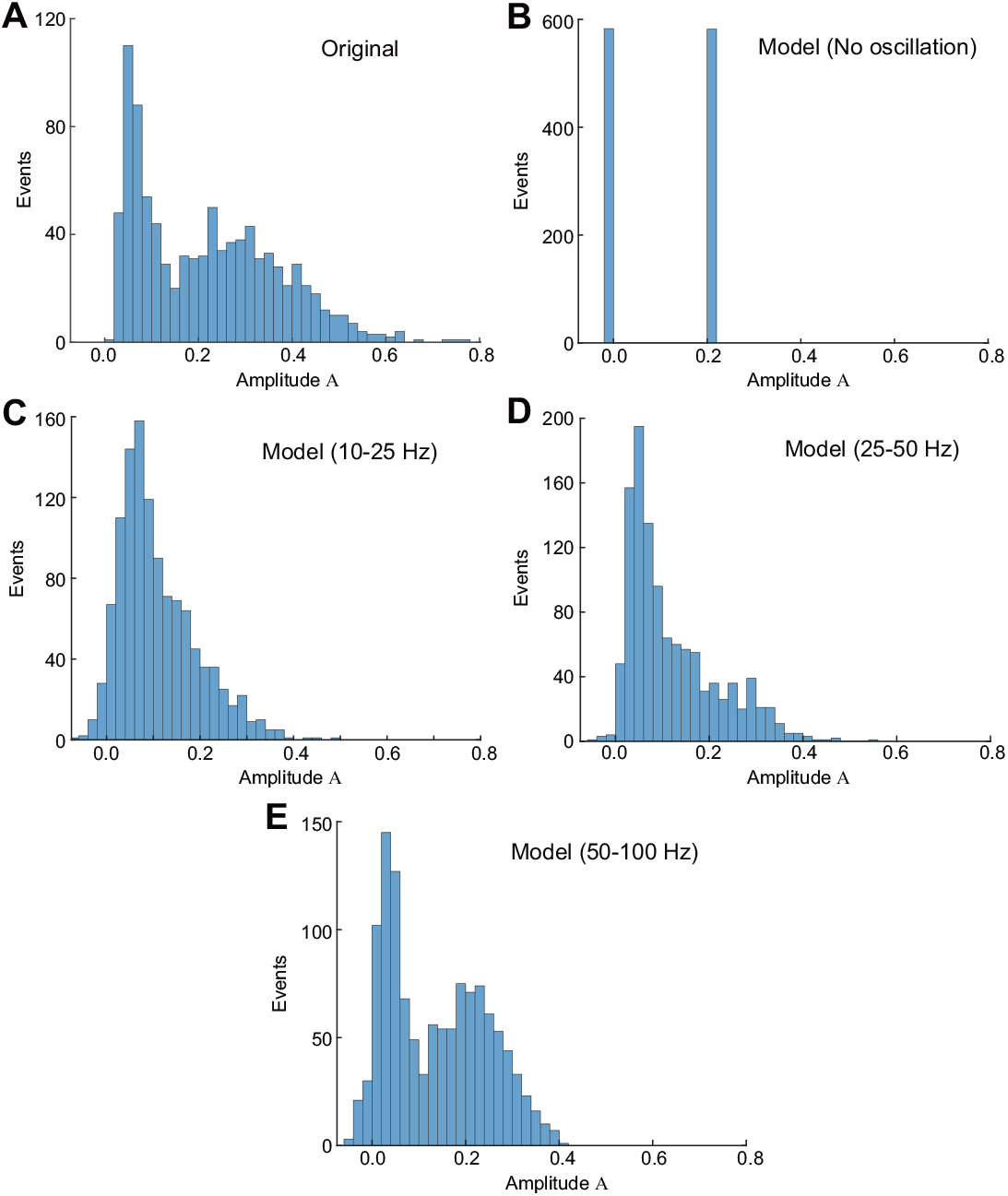
Histograms of hi-hat amplitudes. A: the original performance data. B: the model output without input oscillators. C, D, E: the model outputs with oscillators in the frequency ranges of [10, 25] Hz, [25, 50] Hz, and [50, 100] Hz, respectively.

Second, DFA, a method of studying long-range correlations in time-series data, particularly 1*/f* noise [15], was applied to the timing intervals and amplitudes. Fig 6 shows the results of DFA for the timing interval fluctuations. For the original performance data, the DFA plot can be approximated by two lines bordered by a scale of 40 (Fig 6A). Table 2 lists the corresponding exponents for the shorter and longer scales in the *α*_*τ*_ (short) and *α*_*τ*_ (long) columns. The exponent values for original data were 0.31 (anticorrelations) and 0.72 (correlations). In contrast, the exponent for the model without oscillators was approximately zero (Fig 6B), reflecting that its timing interval series lacked significant fluctuations. The DFA plots for the models with low-frequency oscillators ([10, 25] Hz and [25, 50] Hz) did not show a clear separation between two scales (Fig 6C and D). Their exponent values were approximately 0.5, which indicates that the fluctuations were uncorrelated, similar to white noise. The analysis of the high-frequency model ([50, 100] Hz) output yielded a DFA plot with two distinct slopes, similar to that of the original data (Fig 6E). However, both of its exponents were less than 0.5, indicating the presence of anti-persistent correlations across both short and long scales.

**Fig 6.**
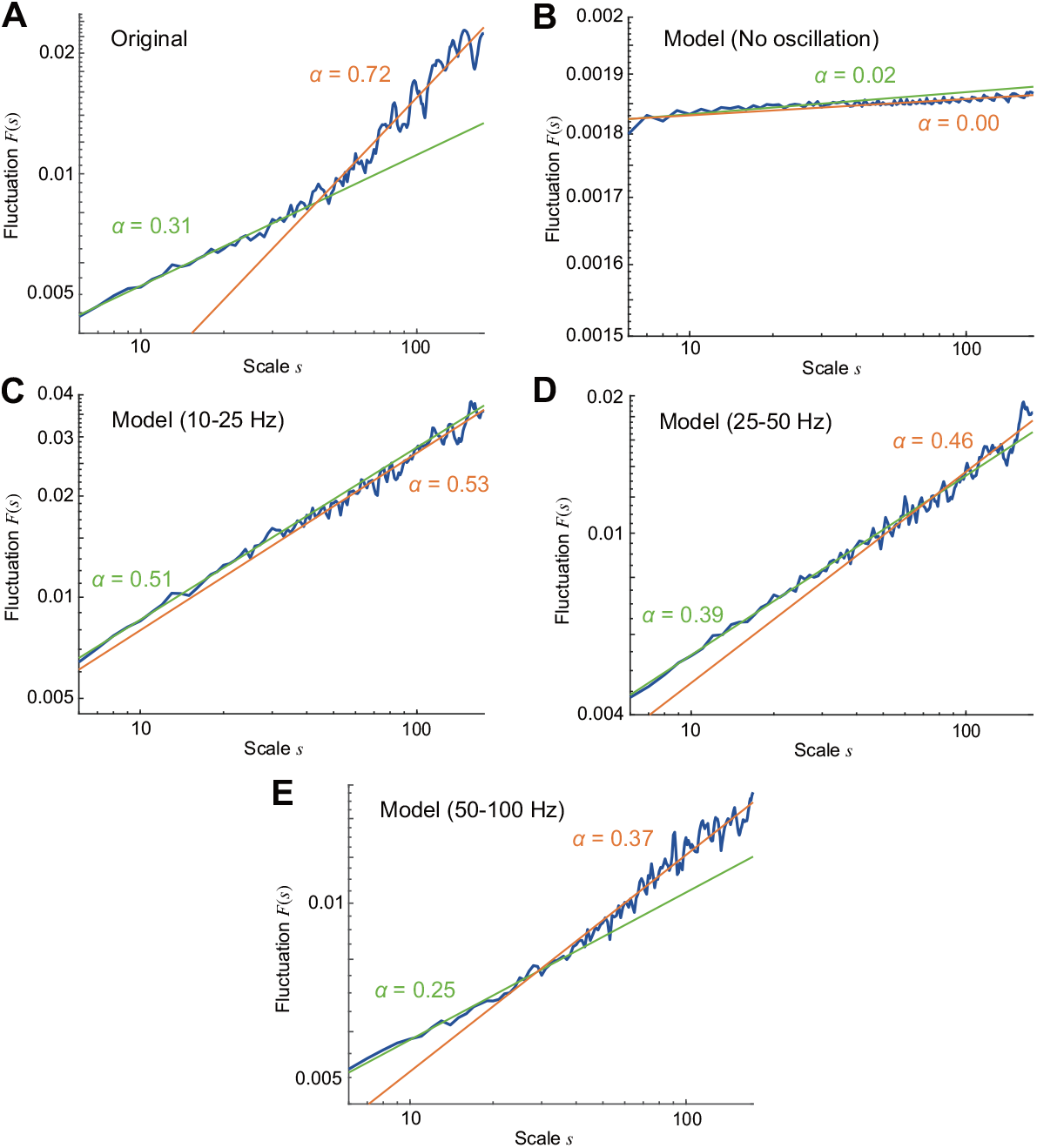
Results from DFA for timing intervals. A: the original performance data. B: the model output without input oscillators. C, D, E: the model outputs with oscillators in the frequency ranges of [10, 25] Hz, [25, 50] Hz, and [50, 100] Hz, respectively.

Unlike the timing intervals, the DFA for amplitude fluctuations exhibited a single, uniform scaling region across all conditions, as shown in Fig 7. The corresponding scaling exponents (*α*_*A*_) are listed in Table 2. The original performance data yielded an exponent greater than 0.5 (*α*_*A*_ *>* 0.5), indicating persistent, long-range correlations (Fig 7A). In contrast, all model outputs that included oscillators produced exponents less than 0.5 (*α*_*A*_ *<* 0.5), which is characteristic of anti-persistent correlations (Fig 7C–E). This anti-persistence suggests a mean-reverting dynamic, where large amplitude values are likely to be followed by small ones, and vice versa.

**Fig 7.**
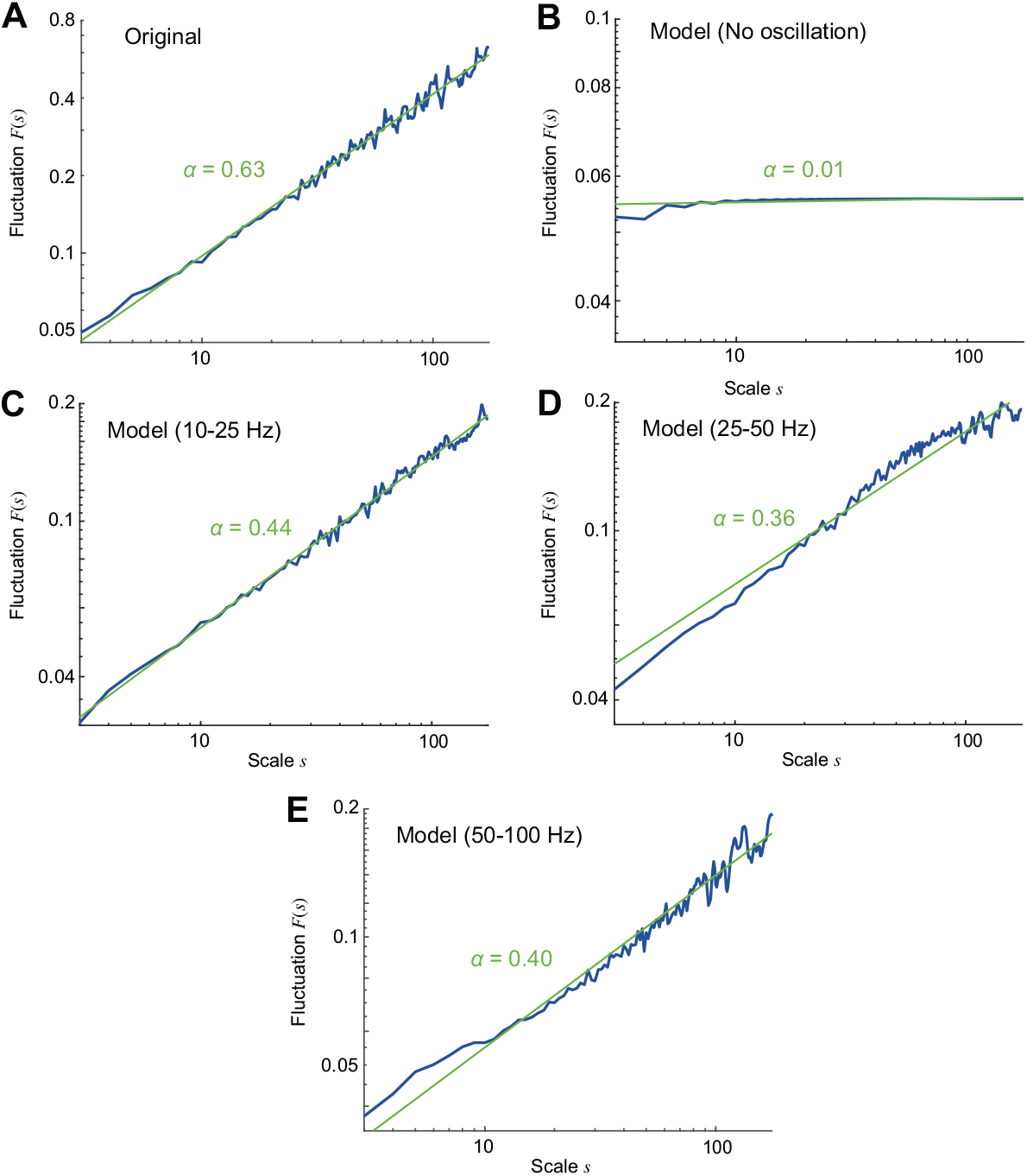
Results from DFA for amplitudes. A: the original performance data. B: the model output without input oscillators. C, D, E: the model outputs with oscillators in the frequency ranges of [10, 25] Hz, [25, 50] Hz, and [50, 100] Hz, respectively.

Third, we analyzed the temporal dynamics of the timing intervals using return maps (Fig 8). These maps are scatter plots created by plotting each timing interval *τ*_*i*_ against the subsequent one (*τ*_*i*+1_). The average correlation coefficients (*r*) from these plots over 20 runs are summarized in Table 2. The return map of the original performance data exhibited a negative correlation (*r* = − 0.48), indicating a tendency for a shorter interval to be followed by a longer one, and vice versa (Fig 8A). For the model without oscillators, the points formed two distinct clusters, revealing an alternating pattern between two specific timing intervals (Fig 8B). In contrast, introducing low-frequency inputs to the model resulted in an uncorrelated plot (Fig 8C). Notably, the correlation became progressively more negative as the input frequencies increased (Fig 8D and E). The model with high-frequency oscillators ([50, 100] Hz) successfully replicated the negative correlation observed in the original data (*r* = −0.48).

**Fig 8.**
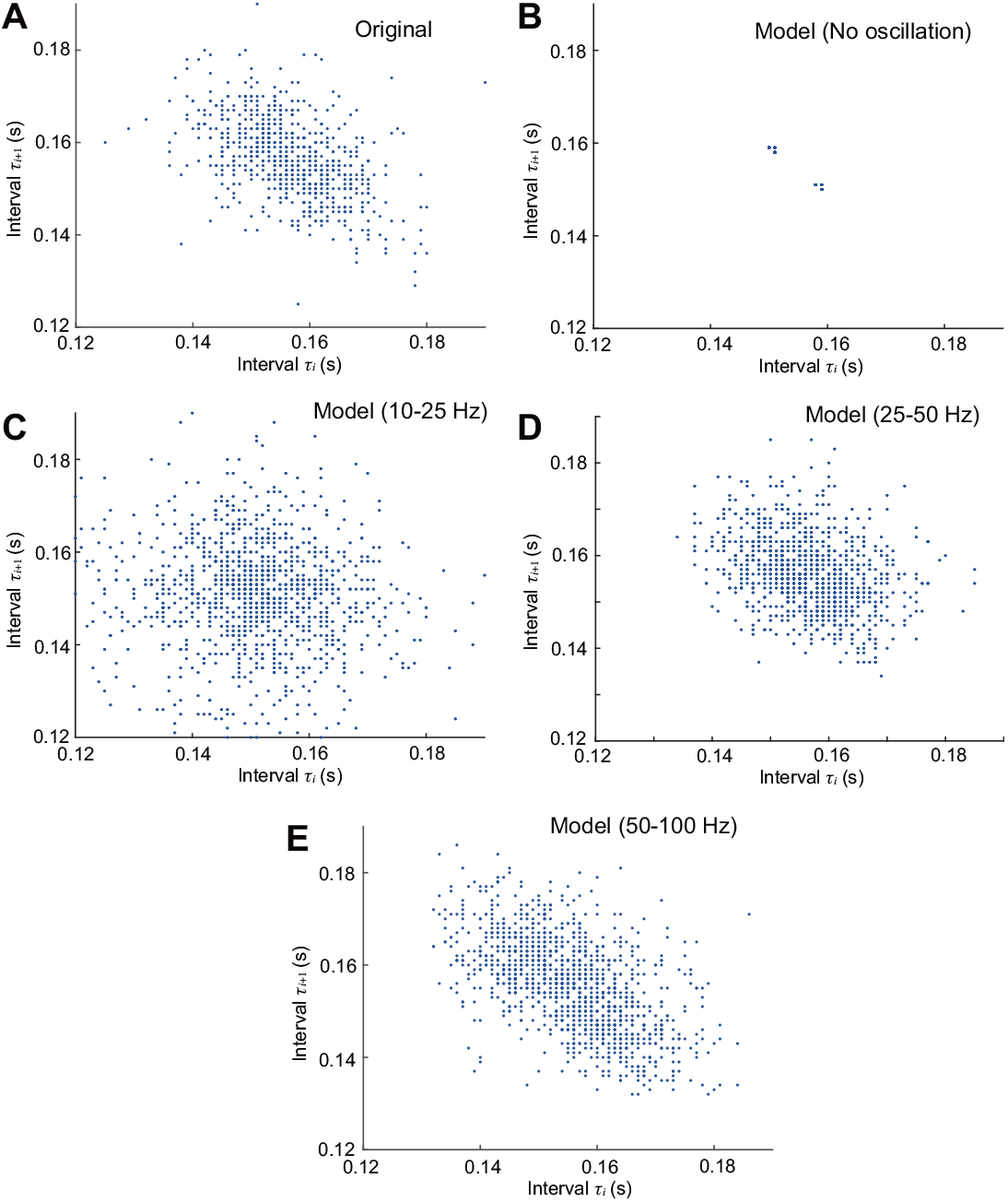
Return map of timing intervals. A: the original performance data. B: the model output without oscillators. C, D, E: the model outputs with oscillators in the frequency ranges of [10, 25] Hz, [25, 50] Hz, and [50, 100] Hz, respectively.

Finally, we analyzed the short-scale temporal patterns within the timing intervals, focusing on segments equivalent to bars consisting of 16 hits. Fig 9A shows the 16th-note intervals from the first 10 bars of the original performance data (the song’s introduction). A “short-long-short” pattern is observed at the beginning of the bars.

**Fig 9.**
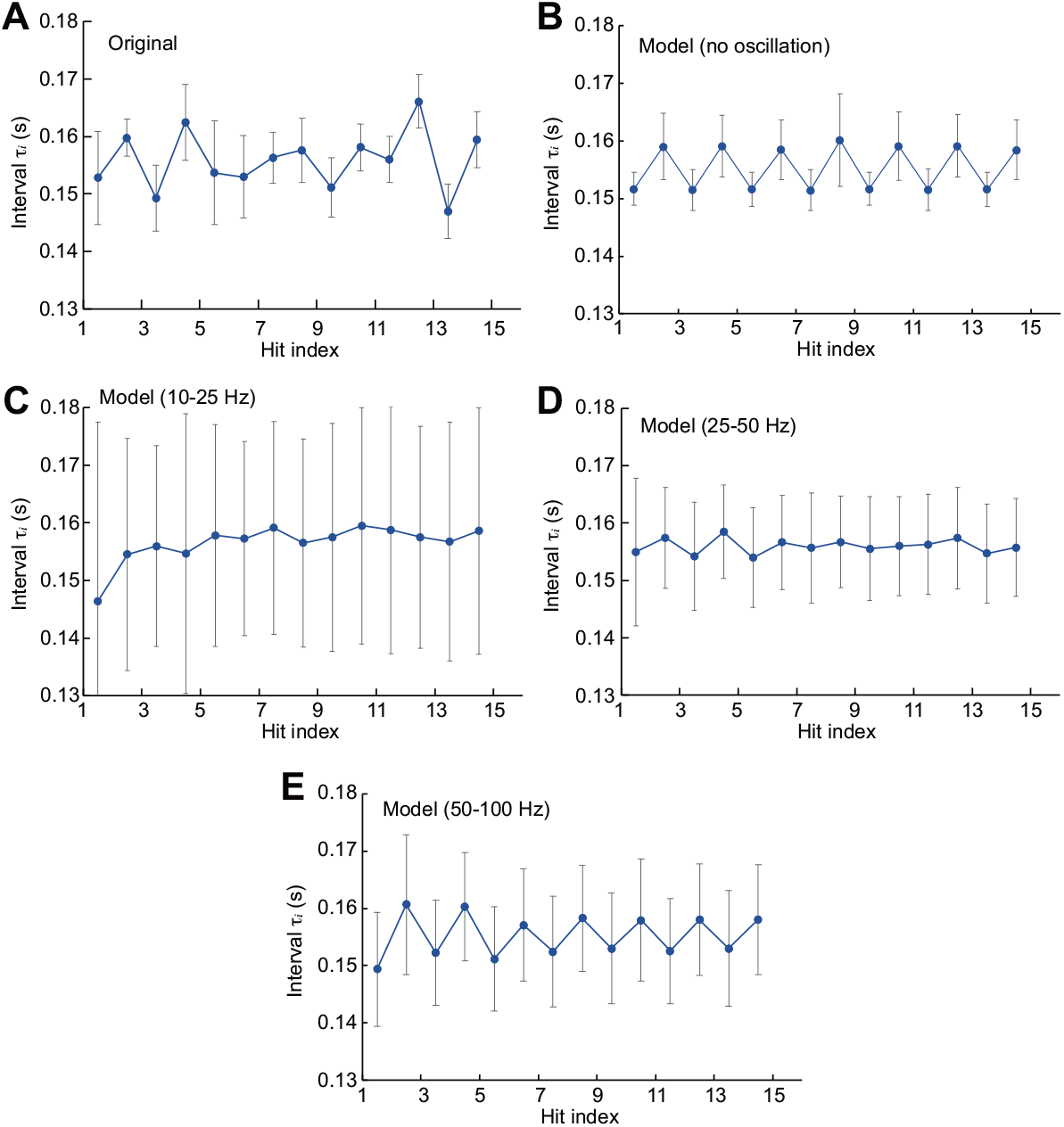
Bar patterns of timing intervals. A: The first 10 bars of the original data. B: the model output without input oscillators. C, D, E: the model outputs with oscillators in the frequency ranges of [10, 25] Hz, [25, 50] Hz, and [50, 100] Hz, respectively. Error bars indicate the standard deviation.

Since the model outputs lacked a predefined bar structure, we applied different segmentation method. For the model without oscillators, we segmented the time series into non-overlapping, consecutive boxes of 16 data points to create artificial “bars,” starting the first box at an initial short timing interval. The averaged pattern across all such boxes, calculated over 20 runs, revealed a “short-long-short” sequence (Fig 9B). For the models with oscillators, we defined bars as segments where a 16th rest was followed by 15 16th notes. The resulting patterns, averaged over 20 runs, showed that the low-frequency models ([10, 25] Hz and [25, 50] Hz) did not produce a clear or consistent pattern (Fig 9C and D). In contrast, the high-frequency model ([50, 100] Hz) exhibited a clear “short-long-short” pattern following the 16th rests (Fig 9E).

A similar analysis was conducted for the amplitude series (Fig 10). The original performance data exhibited a distinct “strong-weak-strong” pattern (Fig 10A). The model without oscillators clearly and consistently produced this “strong-weak-strong” amplitude pattern (Fig 10B). In contrast, the outputs from the low-frequency model ([10, 25] Hz) lacked this structure (Fig 10C). The high-frequency models ([25, 50] Hz and [50, 100] Hz) successfully generated the “strong-weak-strong” pattern (Fig 10D and E). However, the nature of accentuation differed from the original; while the primary accents (on the 1st, 5th, 9th, and 13th hits) were more pronounced in the original pattern, the nuanced contour was not replicated by the model.

**Fig 10.**
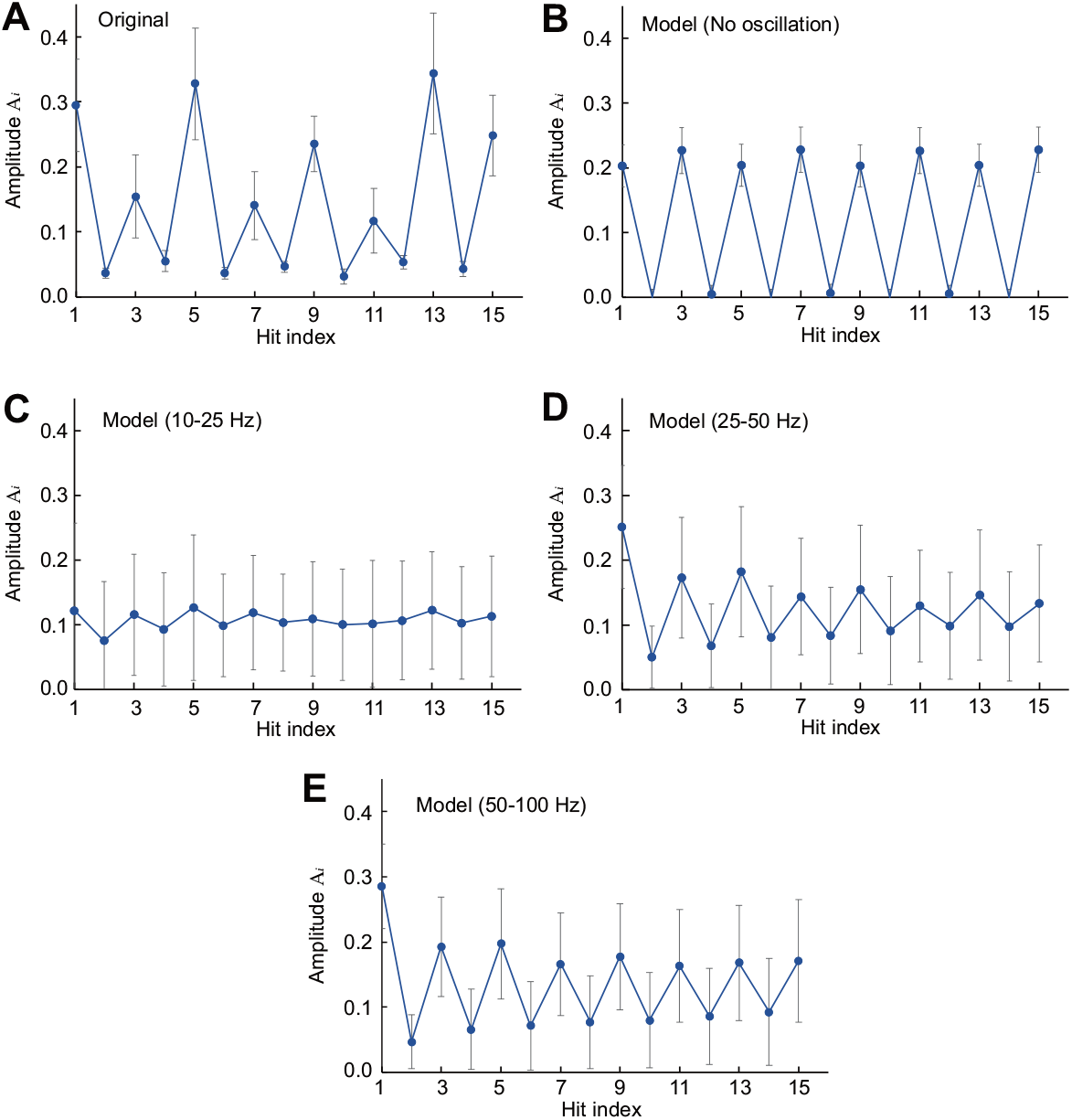
Bar patterns of amplitudes. A: The first 10 bars of the original data. B: the model output without input oscillators. C, D, E: the model outputs with oscillators in the frequency ranges of [10, 25] Hz, [25, 50] Hz, and [50, 100] Hz, respectively. Error bars indicate the standard deviation.

### Learning drum-kit performances

Next, the reservoir computing model with [50, 100]-Hz oscillators was trained using the target encoded from the multidimensional drum-kit performances. From the Groove MIDI Dataset [68], funk, jazz, samba, and rock files were selected and the first 20 s of each was used as a target. The hit onset timings’ and their amplitudes’ data were extracted from the MIDI files and encoded into time-series data, as in the case of the hi-hat detailed above. A readout layer was provided for each multiple-drum instrument. The audio files of the targets are attached as Audios S6 to S9.

Figs 11–14 respectively show example outputs of the model trained with the funk, jazz, samba, and rock performances. Their audio files are attached as Audios S10 to S13. In each case, a performance similar to the target was generated. In the case of funk (Fig 11), a series of snare hits emerged, which was not present. The jazz-learned model reproduced its characteristic fill-in (tom, snare, and kick patterns in that order), indicating the model learned such spatiotemporal pattern (Fig 12). The samba output was relatively periodic, but the snare patterns dynamically transitioned (Fig **??**). In the case of rock (Fig 14), a clear inter-modality rule was acquired that a snare (rim) was not struck when a snare (head) was hit with a large amplitude. These results demonstrate that the model could learn the coordination between drums to create dynamic, complex rhythmic patterns.

**Fig 11.**
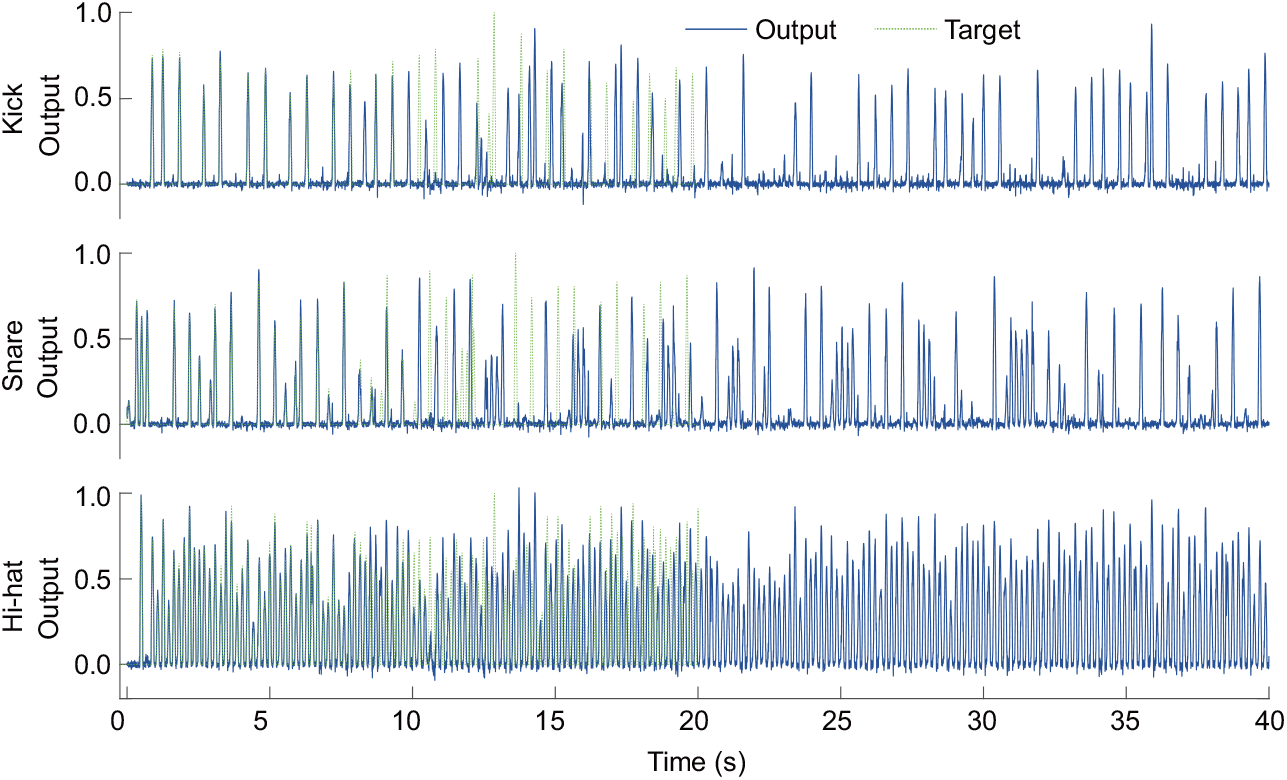
Example outputs of the reservoir model trained with the funk performance. The solid-blue and dashed-green curves respectively represent the model’s output and the target.

**Fig 12.**
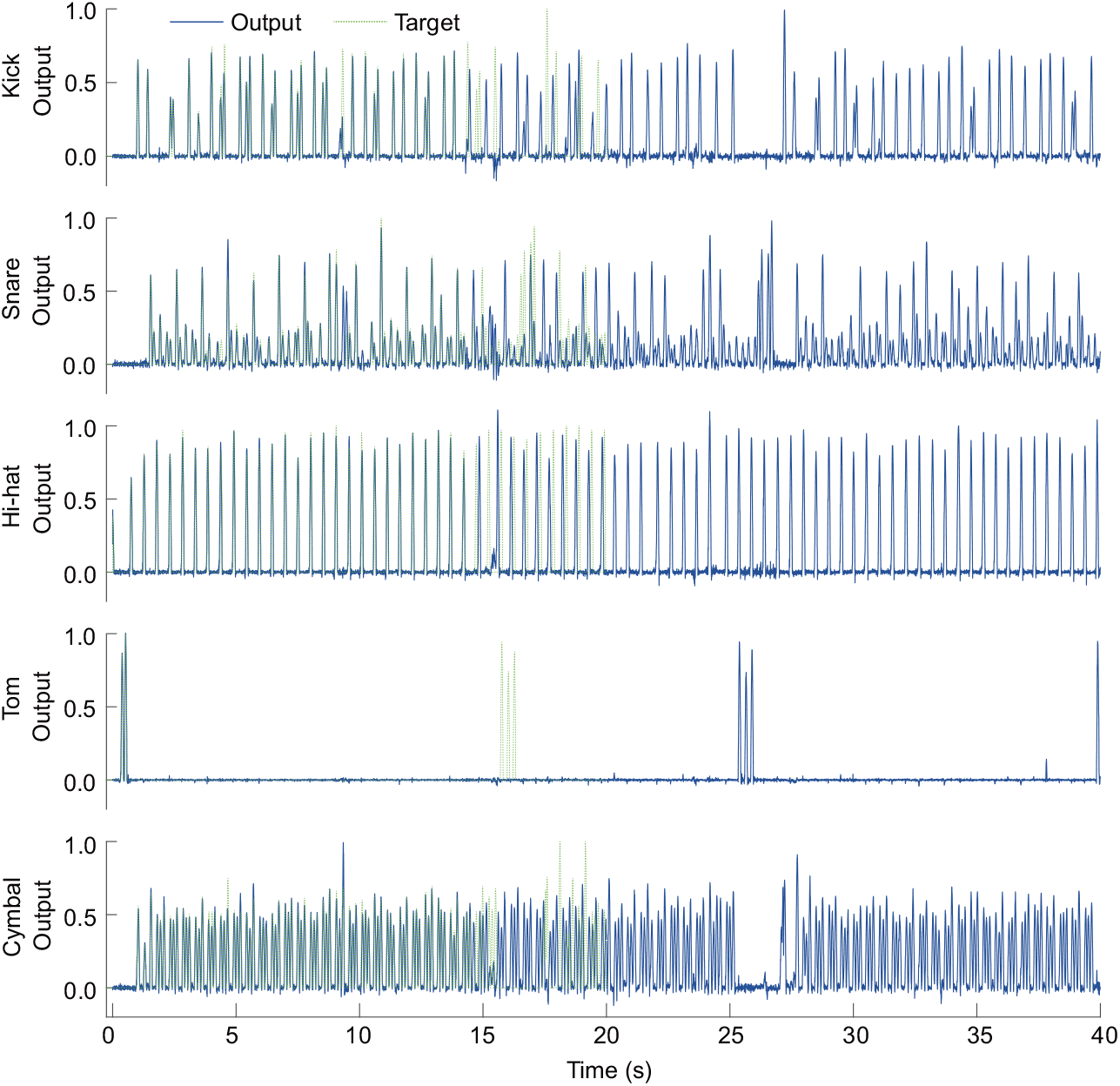
Example outputs of the reservoir model trained with the jazz performance. The solid-blue and dashed-green curves respectively represent the model’s output and the target.

**Fig 13.**
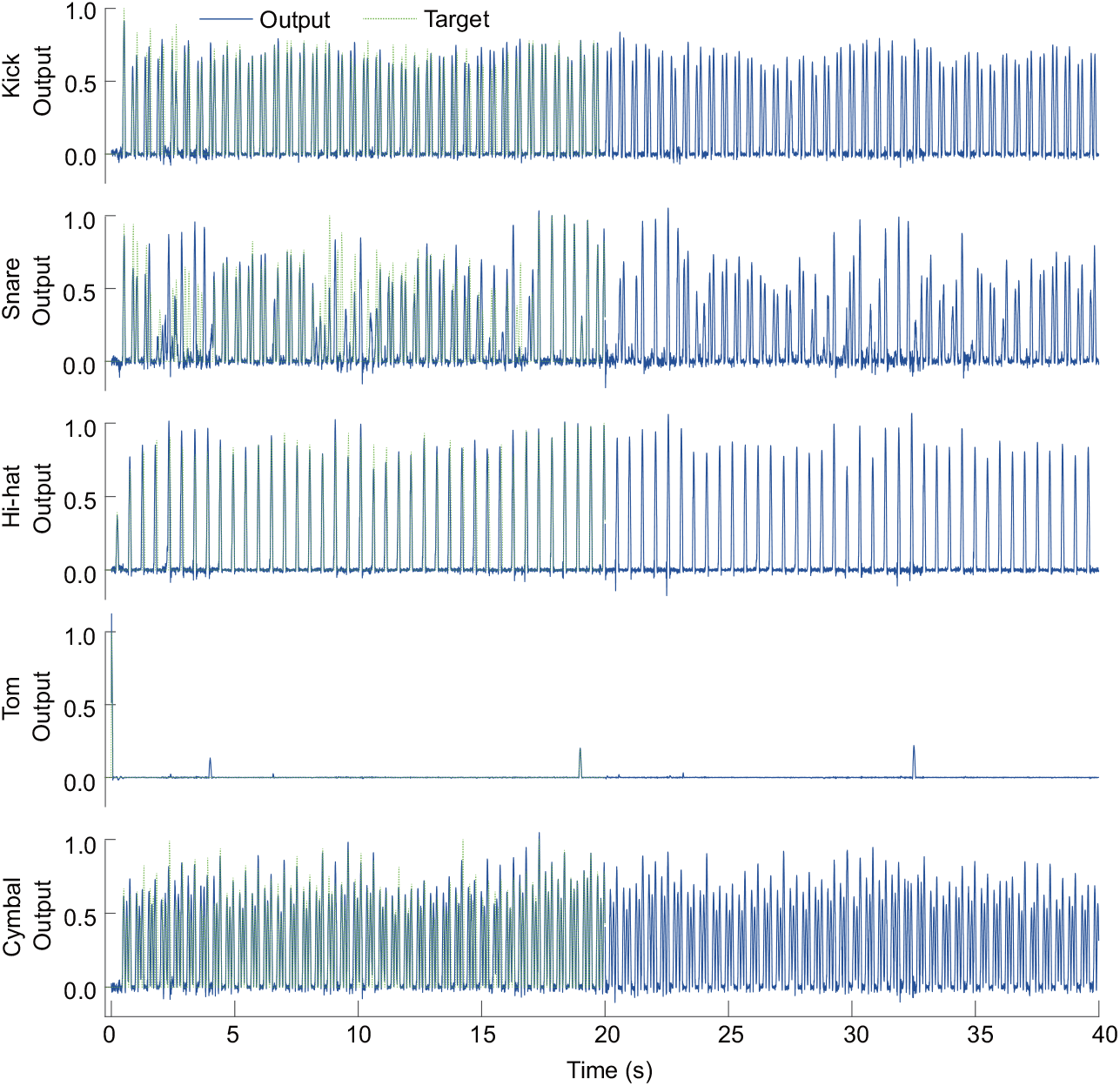
Example outputs of the reservoir model trained with the samba performance. The solid-blue and dashed-green curves respectively represent the model’s output and the target.

**Fig 14.**
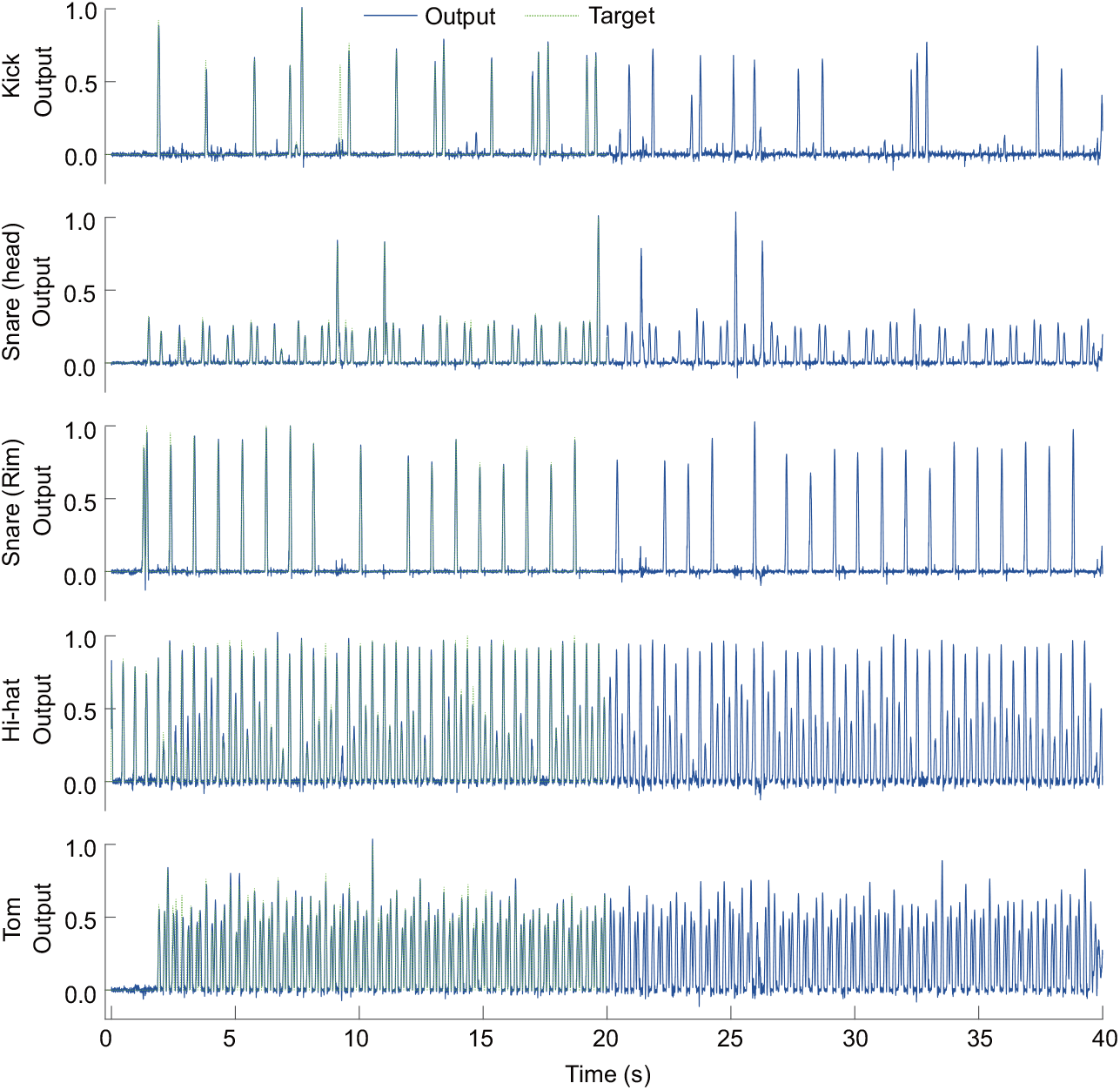
Example outputs of the reservoir model trained with the rock performance. The solid-blue and dashed-green curves respectively represent the model’s output and the target.

We investigated whether microtiming of the hi-hat performances in the model outputs reproduced that of the originals. The hi-hat rhythm was better suited for this microtiming analysis because it was more periodic than those of other drum parts. 20 models with different random seeds each output 60 seconds, and their hi-hat hit timing intervals were calculated. The hi-hat intervals for the entire original performances were also calculated. Fig 15 shows histograms of the hi-hat inter-beat intervals of the four original performances and model outputs. Histograms of the original performances showed bimodal peaks because the original hi-hat rhythm changed in the middle of the performances; therefore, in Fig 15, the original histograms were cut out at the same intervals as the histograms of the model outputs. The means and standard deviations of those histograms are listed in Table 3. These results indicate that the average timing intervals matched between the originals and model outputs, whereas the microtiming variability of the model outputs was greater than that of the originals. This increased variability is a property of the model, as it was also observed in the previous hi-hat performance analysis (see Fig 4 and the *τ* column in Table 2).

**Table 3.** Means and standard deviations of microtiming.

| Genre | Source | Mean (s) | Standard deviation (s) |
| --- | --- | --- | --- |
| Funk | Original | 0.186 | 0.0108 |
|  | Model | 0.188 | 0.0207 |
| Jazz | Original | 0.519 | 0.0115 |
|  | Model | 0.518 | 0.0219 |
| Samba | Original | 0.517 | 0.0144 |
|  | Model | 0.517 | 0.0187 |
| Rock | Original | 0.240 | 0.0194 |
|  | Model | 0.241 | 0.0199 |

**Fig 15.**
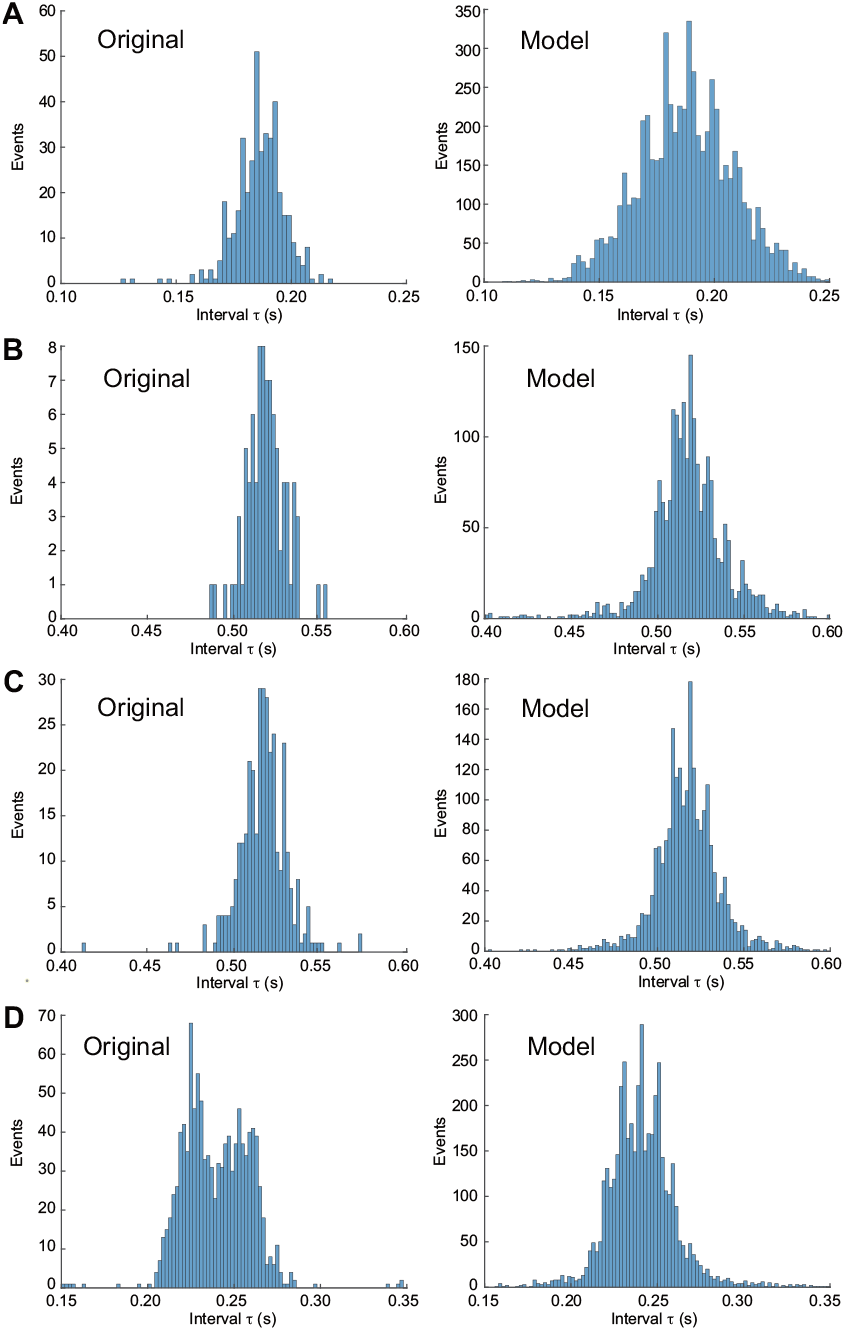
Histograms of the timing intervals of hi-hat hits in the four original performances (left panels) and model outputs (right panels). A–D: timing intervals for the funk, jazz, samba, and rock performances, respectively.

The similarity of the audio features between the original performances and model outputs was quantified. We calculated six audio features related to groove feel [21] from the audio files of the original performances and the 20 model outputs. The values of these audio features are shown in Fig 16. This figure shows that most of the audio features of the originals and model outputs were consistent, indicating their acoustical similarity. For pulse clarity (Fig 16D), the values of the model outputs were smaller than those of the originals. Since pulse clarity quantifies the periodicity of a rhythm, this result indicates that the periodicity or metric-structural components of the model outputs was reduced.

**Fig 16.**
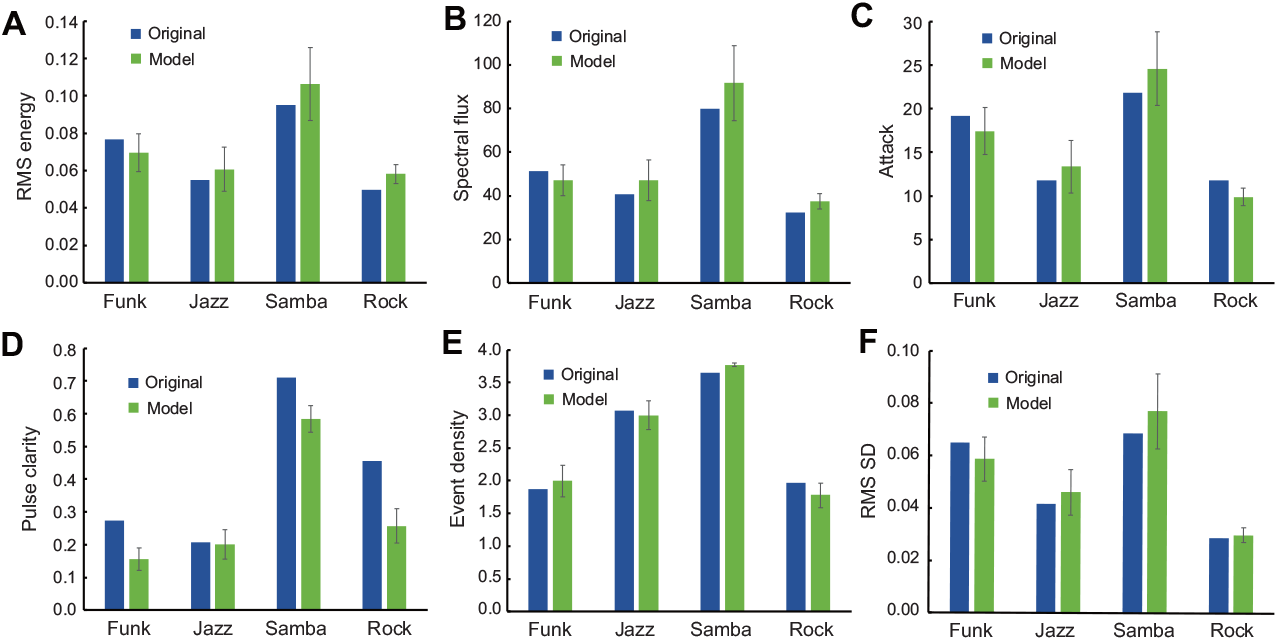
Audio features of the original performances (blue) and the model outputs (green). A–F: bar plots for RMS energy, spectral flux, attack, pulse clarity, event density, RMS SD, respectively. In model outputs, the error bars indicate the standard deviations among 20 models.

## Discussion

We developed a simple RNN model that learns complex drum performances using the oscillation-driven reservoir computing. This model learned real-world drum performances and replicated the fluctuations of inter-beat timings (microtiming) and audio features relating to a sense of groove. Furthermore, it could generalize the short target data samples to generate rhythms that were close to but different from the target, which could be called improvisation based on an example. In the learning task of playing a multidimensional drum kit, the model could learn the spatiotemporal patterns between the instruments, resulting in dynamic and consistent performances, including fill-ins. Most existing models of beat perception and generation directly adjust the frequencies of neural oscillators to fit isochronous target rhythms [59–62]. This approach allows only a simple rhythm to be generated. In the present model, the oscillator frequencies are fixed, and the network dynamics are tuned via readout training, which enables the model to learn complex real-world rhythms.

Oscillation-driven reservoir computing was designed for learning timings [54]. Time can be measured by reservoir activity driven by oscillator inputs, even during periods of no external stimulus. In this study, oscillators were used to generate complex reservoir activity. Conventional reservoir computing without oscillators produces simple periodic outputs. Although these outputs exhibited basic alternating patterns, such as a “short-long-short” pattern in timing interval series (Fig 9B) and a “strong-weak-strong” pattern in amplitude series (Fig 10B), they lacked the fluctuations characteristic of human performance. Consequently, applying analyses such as DFA and return maps to these outputs yielded no informative insights into such dynamics (Figs 6B, 7B, and 8B). In contrast, the introduction of high-frequency oscillators enable the reservoirs to generate human-like complex rhythms that not only exhibit structured patterns like “short-long-short” and “strong-weak-strong” (Figs 9E and 10E) but also possess the kind of long-range correlated fluctuations found in professional human performances (Figs 6E, 7E, and 8E). Therefore, we suggest this as a unified model for learning timings and rhythms. In the human brains, the cerebellum and basal ganglia are thought to be responsible for such learning [1, 20, 22–27]. Indeed, the fact that previous computational models in these areas also used a reservoir computing framework [42–47] further supports our model.

The model’s performance depended on the given frequency bands. Low-frequency ([10, 25] Hz) inputs contributed to the accurate reproductions of the targets, whereas high-frequency ([50, 100] Hz) inputs contributed to generalizing the target to generate target-like performances. This tendency is similar to that observed in previous studies on chaotic time-series prediction [54, 82]. We hypothesize that the low-frequency inputs acted as “time stamps” for the targets. That is, the model learned the relationship between the specific dynamics induced by the oscillator inputs, which represented time, and the targets. However, the model that overfitted the relationship between specific reservoir states and the target was unable to generalize it to unlearned states. In contrast, high-frequency inputs did not significantly affect the reservoir states because the neuron model had a time constant and did not change states at high speed. Therefore, the model somewhat preserved the network-inherent dynamics, and thus prevented overfitting.

This suggests that different neural mechanisms are involved in for copying and improvising musical performances. High-frequency (beta- and gamma-band) oscillations were observed in magnetoencephalography (MEG) during music beat perception, imagery, and timing prediction [83, 84]. An electroencephalogram (EEG) study by Rosen et al. [85] found clusters of beta- and gamma-band activity when higher-quality and lower-quality improvisations were compared in jazz performances. Subsequently, a group of the authors [86] recorded jazz guitarists’ EEGs while improvising to provide chord sequences. They found that high-flow (effortless attention to the task) was associated with high-frequency (gamma-band) clusters in the left opercula and in the left temporal gyri. Such high-frequency cortical activity may facilitate the generalization of musical performances. Using EEGs to detect further high-frequency (high-gamma-band) signals is difficult, but studies using electrocorticography (ECoG) have revealed that rhythmic information is contained in the high-gamma signals in the auditory cortex [87–89]. This confirms that listening to music induces high-frequency signals, and it would be interesting to study how this relates to improvising performances.

Whatever frequency bands were used, these were not directly related to the tasks of the rhythm learning. For example, the average inter-beat intervals for the hi-hat performance is 0.156 s, which corresponds to approximately 6.4 Hz. The frequencies of the input oscillators were much higher than those and were randomly determined. Even though the model did not have an external “metronome,” it acquired an internal representation of a metronome as a result of training, resulting in stable performances.

In learning the hi-hat rhythm, the model generated performances with fluctuations in the timing intervals and amplitudes. These fluctuations were not random but exhibited the same distributions and patterns (e.g., the high-low-high-low pattern) as the original fluctuations. Such repetitive timing deviation is called *systematic* microtiming [17, 90], and microtiming within the same instrument is considered *horizontal*. Therefore, the model could copy *systematic horizontal* microtiming of the professional hi-hat performance. However, the long-range correlations of the model fluctuations were negative, showing an inverse relationship to those of the original fluctuations. This means that the model outputs were stable; that is, an increase at one time point tended to be followed by a decrease, and vice versa. In addition, this model could not learn the concepts of bars and meters. The 16th rests, which marks the end of the bars in the original hi-hat performance, was randomly placed in the model’s outputs. Therefore, this model fails to learn the long-term rhythmic patterns. A solution may conceivably be to include meter information in the oscillators. An EEG study revealed that gamma-band signals in the auditory cortex reflect the metric structure [91]. Other studies have also shown that brain oscillations in various frequency bands contain information about the meters of rhythms being heard [83, 92, 93]. By including meter information in the oscillations, it may be possible to learn long-term rhythmic patterns.

In learning the drum-kit performances of various genres, the model generated outputs similar to the original performances. The microtiming analysis of the hi-hat performances showed that the model outputs had horizontal microtiming with greater variability than the originals. This might be due to the variability in neural dynamics of the model. Nevertheless, their standard deviations were less than 20 ms, which are considered as microtiming, defined as timing deviation of 50 ms or less. The audio feature analysis showed that with except for the pulse clarity feature, the model outputs copied the features that correlate with a sense of groove, indicating that the model can copy groovy performances. The pulse clarity values for periodicity were reduced because the model did not represent long-term structures, including meter and bar, as discussed earlier.

This model uses professional performance as a target and copies it in just ten training sessions. Therefore, it does not explain how the drumming performance improves. In reality, performance skills are acquired gradually during numerous practice routines designed and arranged by teachers and coaches [94, 95]. Performing music involves more than a single neural circuit but a variety of sensorimotor and cognitive functions [96], such as motor control [97, 98], planning and temporal control [99], and feedback monitoring [100]. An important future research direction is to investigate the processes for acquiring such complex skills to construct their computational models.

## Conclusion

In this study, we investigated the capacity of an oscillation-driven reservoir computing model to learn and generate complex drum rhythms. Our findings demonstrate that the model with high-frequency inputs can successfully acquire and reproduce the subtle microtiming variations and expressive dynamics that are characteristic of professional drumming. This suggests that a relatively simple RNN architecture is sufficient to capture the fine-grained temporal patterns.

However, we identified a notable limitation: the model was unable to learn higher-level rhythmic structures, such as meter and bar divisions. While it could generate locally coherent patterns, it failed to adhere to the long-term, hierarchical organization that governs most musical rhythms. This distinction suggests that the cognitive and computational mechanisms for processing local, expressive timing may be distinct from those required for understanding abstract, metrical hierarchies. Future research should focus on developing models that can integrate these two levels of rhythmic processing. Such work will be crucial for building a more comprehensive computational theory of human rhythmic capability.

## Supporting information

- Audio S1. Audio file of the hi-hat learning target.
- Audio S2. Audio file of an output of the model without oscillations.
- Audio S3. Audio file of an output of the model with [10, 25]-Hz oscillators.
- Audio S4. Audio file of an output of the model with [25, 50]-Hz oscillators.
- Audio S5. Audio file of an output of the model with [50, 100]-Hz oscillators.
- Audio S6. Audio file of the funk target.
- Audio S7. Audio file of the jazz target.
- Audio S8. Audio file of the samba target.
- Audio S9. Audio file of the rock target.
- Audio S10. Audio file of a model output trained with the funk.
- Audio S11. Audio file of a model output trained with the jazz.
- Audio S12. Audio file of a model output trained with the samba.
- Audio S13. Audio file of a model output trained with the rock.

---

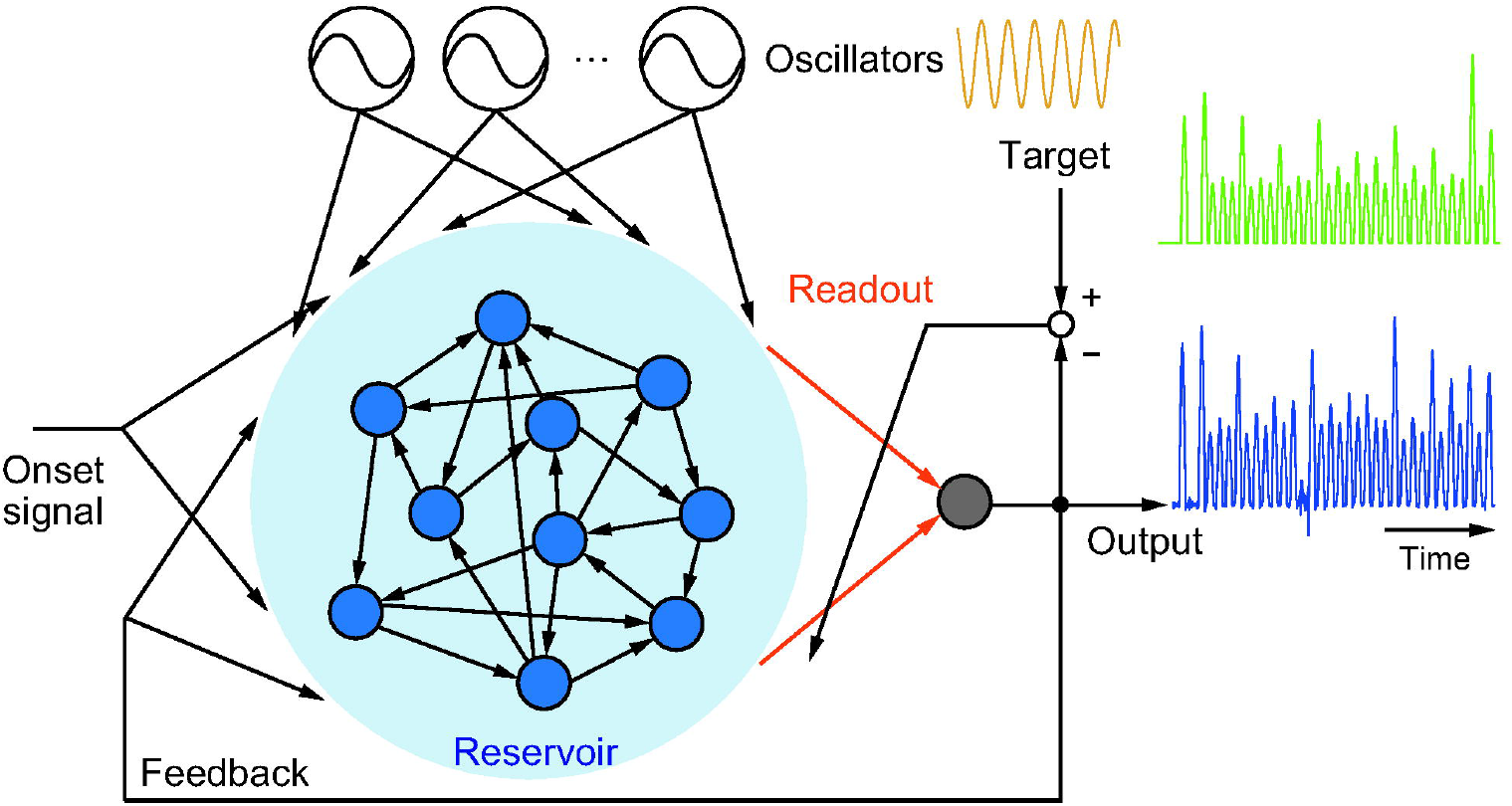

## References

1. Grahn JA. Neural mechanisms of rhythm perception: current findings and future perspectives. Topics in Cognitive Science. 2012;4(4):585–606.

2. London J. Hearing in Time: Psychological Aspects of Musical Meter. Oxford University Press; 2012.

3. Patel AD, Iversen JR. The evolutionary neuroscience of musical beat perception: the Action Simulation for Auditory Prediction (ASAP) hypothesis. Frontiers in Systems Neuroscience. 2014;8:57.

4. Large EW, Roman I, Kim JC, Cannon J, Pazdera JK, Trainor LJ, et al. Dynamic models for musical rhythm perception and coordination. Frontiers in Computational Neuroscience. 2023;17:1151895.

5. Povel DJ. Internal representation of simple temporal patterns. Journal of Experimental Psychology: Human Perception and Performance. 1981;7(1):3–18.

6. Iyer V. Embodied mind, situated cognition, and expressive microtiming in African-American music. Music perception. 2002;19(3):387–414.

7. Fujii S, Hirashima M, Kudo K, Ohtsuki T, Nakamura Y, Oda S. Synchronization error of drum kit playing with a metronome at different tempi by professional drummers. Music Perception: An Interdisciplinary Journal. 2011;28(5):491–503.

8. Räänen E, Pulkkinen O, Virtanen T, Zollner M, Hennig H. Fluctuations of hi-hat timing and dynamics in a virtuoso drum track of a popular music recording. PLOS One. 2015;10(6):e0127902.

9. Câmara GS, Nymoen K, Lartillot O, Danielsen A. Timing is everything… or is it? Effects of instructed timing style, reference, and pattern on drum kit sound in groove-based performance. Music Perception: An Interdisciplinary Journal. 2020;38(1):1–26.

10. Nelias C, Sturm EM, Albrecht T, Hagmayer Y, Geisel T. Downbeat delays are a key component of swing in jazz. Communications Physics. 2022;5(1):237.

11. Kilchenmann L, Senn O. Microtiming in swing and funk affects the body movement behavior of music expert listeners. Frontiers in psychology. 2015;6:1232.

12. Iyer VS. Microstructures of feel, macrostructures of sound: Embodied cognition in West African and African-American musics. University of California, Berkeley; 1998.

13. Butterfield MW. The Power of Anacrusis: Engendered Feeling in Groove-Based Musics. Music Theory Online. 2006;12(4).

14. McDonald M. In: I Keep Forgettin’. Warner Bros; 1982..

15. Peng CK, Havlin S, Stanley HE, Goldberger AL. Quantification of scaling exponents and crossover phenomena in nonstationary heartbeat time series. Chaos: An Interdisciplinary Journal of Nonlinear Science. 1995;5(1):82–7.

16. Madison G. Experiencing groove induced by music: consistency and phenomenology. Music Perception. 2006;24(2):201–8.

17. Madison G, Gouyon F, Ullén F, Hörnström K. Modeling the tendency for music to induce movement in humans: first correlations with low-level audio descriptors across music genres. Journal of Experimental Psychology: Human Perception and Performance. 2011;37(5):1578–94.

18. Janata P, Tomic ST, Haberman JM. Sensorimotor coupling in music and the psychology of the groove. Journal of Experimental Psychology: General. 2012;141(1):54.

19. Witek MAG, Clarke EF, Wallentin M, Kringelbach ML, Vuust P. Syncopation, body-movement and pleasure in groove music. PLOS One. 2014;9(4):e94446.

20. Etani T, Miura A, Kawase S, Fujii S, Keller PE, Vuust P, et al. A review of psychological and neuroscientific research on musical groove. Neuroscience & Biobehavioral Reviews. 2024;158:105522.

21. Stupacher J, Hove MJ, Janata P. Audio features underlying perceived groove and sensorimotor synchronization in music. Music Perception: An Interdisciplinary Journal. 2016;33(5):571–89.

22. Zatorre RJ, Chen JL, Penhune VB. When the brain plays music: auditory–motor interactions in music perception and production. Nature Reviews Neuroscience. 2007;8(7):547–58.

23. Grahn JA, Rowe JB. Feeling the beat: premotor and striatal interactions in musicians and nonmusicians during beat perception. Journal of Neuroscience. 2009;29(23):7540–8.

24. Teki S, Grube M, Griffiths TD. A unified model of time perception accounts for duration-based and beat-based timing mechanisms. Frontiers in Integrative Neuroscience. 2012;5:90.

25. Fujii S, Wan CY. The role of rhythm in speech and language rehabilitation: The SEP hypothesis. Frontiers in Human Neuroscience. 2014;8:777.

26. Merchant H, Grahn J, Trainor L, Rohrmeier M, Fitch WT. Finding the beat: a neural perspective across humans and non-human primates. Philosophical Transactions of the Royal Society B: Biological Sciences. 2015;370(1664):20140093.

27. Kasdan AV, Burgess AN, Pizzagalli F, Scartozzi A, Chern A, Kotz SA, et al. Identifying a brain network for musical rhythm: A functional neuroimaging meta-analysis and systematic review. Neuroscience & Biobehavioral Reviews. 2022;136:104588.

28. Konoike N, Nakamura K. Cerebral substrates for controlling rhythmic movements. Brain Sciences. 2020;10(8):514.

29. Mayville JM, Jantzen KJ, Fuchs A, Steinberg FL, Kelso JAS. Cortical and subcortical networks underlying syncopated and synchronized coordination revealed using fMRI. Human Brain Mapping. 2002;17(4):214–29.

30. Nozaradan S, Schwartze M, Obermeier C, Kotz SA. Specific contributions of basal ganglia and cerebellum to the neural tracking of rhythm. Cortex. 2017;95:156–68.

31. Penhune VB. In: Thaut MH, Hodges D, editors. Musical expertise and brain structure: the causes and consequences of training. Oxford University Press; 2019. p. 419–38.

32. Jueptner M, Weiller C. A review of differences between basal ganglia and cerebellar control of movements as revealed by functional imaging studies. Brain: A Journal of Neurology. 1998;121(8):1437–49.

33. Doya K. Complementary roles of basal ganglia and cerebellum in learning and motor control. Current Opinion in Neurobiology. 2000;10(6):732–9.

34. Ito M. Mechanisms of motor learning in the cerebellum. Brain Research. 2000;886(1-2):237–45.

35. Groenewegen HJ. The basal ganglia and motor control. Neural Plasticity. 2003;10(1-2):107–20.

36. Buhusi CV, Meck WH. What makes us tick? Functional and neural mechanisms of interval timing. Nature Reviews Neuroscience. 2005;6(10):755–65.

37. Grondin S. Timing and time perception: A review of recent behavioral and neuroscience findings and theoretical directions. Attention, Perception, & Psychophysics. 2010;72(3):561–82.

38. Coull JT, Cheng RK, Meck WH. Neuroanatomical and neurochemical substrates of timing. Neuropsychopharmacology. 2011;36(1):3–25.

39. Dreher JC, Grafman J. The roles of the cerebellum and basal ganglia in timing and error prediction. European Journal of Neuroscience. 2002;16(8):1609–19.

40. Teki S, Grube M, Kumar S, Griffiths TD. Distinct neural substrates of duration-based and beat-based auditory timing. Journal of Neuroscience. 2011;31(10):3805–12.

41. Kameda M, Niikawa K, Uematsu A, Tanaka M. Sensory and motor representations of internalized rhythms in the cerebellum and basal ganglia. Proceedings of the National Academy of Sciences. 2023;120(24):e2221641120.

42. Yamazaki T, Tanaka S. The cerebellum as a liquid state machine. Neural Networks. 2007;20(3):290–7.

43. Rössert C, Dean P, Porrill J. At the edge of chaos: how cerebellar granular layer network dynamics can provide the basis for temporal filters. PLOS Computational Biology. 2015;11(10):e1004515.

44. Tokuda K, Fujiwara N, Sudo A, Katori Y. Chaos may enhance expressivity in cerebellar granular layer. Neural Networks. 2021;136:72–86.

45. Dominey PF. Complex sensory-motor sequence learning based on recurrent state representation and reinforcement learning. Biological Cybernetics. 1995;73(3):265–74.

46. Hinaut X, Dominey PF. Real-time parallel processing of grammatical structure in the fronto-striatal system: A recurrent network simulation study using reservoir computing. PLOS One. 2013;8(2):e52946.

47. Kawai Y, Asada M. Spatiotemporal motor learning with reward-modulated Hebbian plasticity in modular reservoir computing. Neurocomputing. 2023;558:126740.

48. Jaeger H. The “echo state” approach to analysing and training recurrent neural networks. GMD-148, German National Research Center for Information Technology. 2001.

49. Jaeger H, Haas H. Harnessing nonlinearity: predicting chaotic systems and saving energy in wireless communication. Science. 2004;304(5667):78–80.

50. Laje R, Buonomano DV. Robust timing and motor patterns by taming chaos in recurrent neural networks. Nature Neuroscience. 2013;16(7):925–33.

51. Vincent-Lamarre P, Lajoie G, Thivierge JP. Driving reservoir models with oscillations: a solution to the extreme structural sensitivity of chaotic networks. Journal of Computational Neuroscience. 2016;41:305–22.

52. Kawai Y, Park J, Tsuda I, Asada M. Learning long-term motor timing/patterns on an orthogonal basis in random neural networks. Neural Networks. 2023;163:298–311.

53. Kawai Y, Park J, Asada M. Reservoir computing using self-sustained oscillations in a locally connected neural network. Scientific Reports. 2023;13(1):15532.

54. Kawai Y, Morita T, Park J, Asada M. Oscillations enhance time-series prediction in reservoir computing with feedback. Neurocomputing. 2025;648:130728.

55. Kawai Y, Morita T, Park J, Asada M. Oscillation-driven reservoir computing for long-term replication of chaotic time series. In: Artificial Neural Networks and Machine Learning – ICANN 2024; 2024. p. 129–41.

56. Large EW, Jones MR. The dynamics of attending: How people track time-varying events. Psychological Review. 1999;106(1):119–59.

57. Large EW, Almonte FV, Velasco MJ. A canonical model for gradient frequency neural networks. Physica D: Nonlinear Phenomena. 2010;239(12):905–11.

58. Large EW, Herrera JA, Velasco MJ. Neural networks for beat perception in musical rhythm. Frontiers in Systems Neuroscience. 2015;9:00159.

59. Mates J. A model of synchronization of motor acts to a stimulus sequence: II. Stability analysis, error estimation and simulations. Biological Cybernetics. 1994;70(5):475–84.

60. Bose A, Byrne Á, Rinzel J. A neuromechanistic model for rhythmic beat generation. PLoS Computational Biology. 2019;15(5):e1006450.

61. Egger SW, L. NM, Jazayeri M. A neural circuit model for human sensorimotor timing. Nature Communications. 2020;11(1):3933.

62. Zemlianova K, Bose A, Rinzel J. A biophysical counting mechanism for keeping time. Biological Cybernetics. 2022;116:205–18.

63. Briot JP, Pachet F. Deep learning for music generation: challenges and directions. Neural Computing and Applications. 2020;32(4):981–93.

64. Ji S, Yang X, Luo J. A survey on deep learning for symbolic music generation: Representations, algorithms, evaluations, and challenges. ACM Computing Surveys. 2023;56(1):1–39.

65. Hutchings P. Talking Drums: Generating drum grooves with neural networks. In: Proceedings of the First International Workshop on Deep Learning and Music joint with IJCNN; 2017. p. 43–7.

66. Makris D, Kaliakatsos-Papakostas M, Karydis I, Kermanidis KL. Conditional neural sequence learners for generating drums’ rhythms. Neural Computing and Applications. 2019;31(6):1793–804.

67. Kingma DP, Welling M. Auto-encoding variational bayes. In: Proceedings of the International Conference on Learning Representations; 2014..

68. Gillick J, Roberts A, Engel J, Eck D, Bamman D. Learning to groove with inverse sequence transformations. In: Proceedings of the 36th International Conference on Machine Learning; 2019..

69. Vaswani A, Shazeer N, Parmar N, Uszkoreit J, Jones L, Gomez AN, et al. Attention is all you need. Advances in Neural Information Processing Systems. 2017;30.

70. Jin C, Wang T, Liu S, Tie Y, Li J, Li X, et al. A transformer-based model for multi-track music generation. International Journal of Multimedia Data Engineering and Management. 2020;11(3):36–54.

71. Huang YS, Yang YH. Pop music transformer: Beat-based modeling and generation of expressive pop piano compositions. In: Proceedings of the 28th ACM International Conference on Multimedia; 2020. p. 1180–8.

72. Nuttall T, Haki B, Jorda S. Transformer neural networks for automated rhythm generation. In: Proceedings of the International Conference on New Interfaces for Musical Expression; 2021..

73. Haykin S. Neural Networks and Learning Machines. 3rd ed. Upper Saddle River, NJ, USA: Pearson; 2009.

74. Pathak J, Hunt B, Girvan M, Lu Z, Ott E. Model-free prediction of large spatiotemporally chaotic systems from data: A reservoir computing approach. Physical Review Letters. 2018;120(2):024102.

75. Chattopadhyay A, Hassanzadeh P, Subramanian D. Data-driven predictions of a multiscale Lorenz 96 chaotic system using machine-learning methods: Reservoir computing, artificial neural network, and long short-term memory network. Nonlinear Processes in Geophysics. 2020;27(3):373–89.

76. Vlachas PR, Pathak J, Hunt BR, Sapsis TP, Girvan M, Ott E, et al. Backpropagation algorithms and reservoir computing in recurrent neural networks for the forecasting of complex spatiotemporal dynamics. Neural Networks. 2020;126:191–217.

77. Linkenkaer-Hansen K, Nikouline VV, Palva JM, Ilmoniemi RJ. Long-range temporal correlations and scaling behavior in human brain oscillations. Journal of Neuroscience. 2001;21(4):1370–7.

78. Parish LM, Worrell GA, Cranstoun SD, Stead SM, Pennell P, Litt B. Long-range temporal correlations in epileptogenic and non-epileptogenic human hippocampus. Neuroscience. 2004;125(4):1069–76.

79. Hennig H, Fleischmann R, Fredebohm A, Hagmayer Y, Nagler J, Witt A, et al. The nature and perception of fluctuations in human musical rhythms. PLOS One. 2011;6(10):e26457.

80. Lartillot O, Toiviainen P. A Matlab toolbox for musical feature extraction from audio. In: the 10th International Conference on Digital Audio Effects; 2007. p. 237–44.

81. Jeff Porcaro;. https://en.wikipedia.org/wiki/Jeff_Porcaro.

82. Park J, Kawai Y, Asada M. Frequency specific effects of oscillatory inputs on timing and chaotic time-series learning in spiking reservoir computing. In: Neural Information Processing (ICONIP 2025); 2025. p. 514–28.

83. Fujioka T, Trainor LJ, Large EW, Ross B. Beta and gamma rhythms in human auditory cortex during musical beat processing. Annals of the New York Academy of Sciences. 2009;1169(1):89–92.

84. Fujioka T, Ross B, Trainor LJ. Beta-band oscillations represent auditory beat and its metrical hierarchy in perception and imagery. Journal of Neuroscience. 2015;35(45):15187–98.

85. Rosen DS, Oh Y, Erickson B, Zhang FZ, Kim YE, Kounios J. Dual-process contributions to creativity in jazz improvisations: An SPM-EEG study. NeuroImage. 2020;213:116632.

86. Rosen D, Oh Y, Chesebrough C, Zhang FZ, Kounios J. Creative flow as optimized processing: Evidence from brain oscillations during jazz improvisations by expert and non-expert musicians. Neuropsychologia. 2024:108824.

87. Sturm I, Blankertz B, Potes C, Schalk G, Curio G. ECoG high gamma activity reveals distinct cortical representations of lyrics passages, harmonic and timbre-related changes in a rock song. Frontiers in Human Neuroscience. 2014;8:798.

88. Ding Y, Zhang Y, Zhou W, Ling Z, Huang J, Hong B, et al. Neural correlates of music listening and recall in the human brain. Journal of Neuroscience. 2019;39(41):8112–23.

89. Herff SA, Herff C, Milne AJ, Johnson GD, Shih JJ, Krusienski DJ. Prefrontal High Gamma in ECoG tags periodicity of musical rhythms in perception and imagination. eNeuro. 2020;7(4).

90. Madison G, Sioros G. What musicians do to induce the sensation of groove in simple and complex melodies, and how listeners perceive it. Frontiers in Psychology. 2014;5:894.

91. Snyder JS, Large EW. Gamma-band activity reflects the metric structure of rhythmic tone sequences. Cognitive Brain Research. 2005;24(1):117–26.

92. Iversen JR, Repp BH, Patel AD. Top-down control of rhythm perception modulates early auditory responses. Annals of the New York Academy of Sciences. 2009;1169(1):58–73.

93. Nozaradan S, Peretz I, Missal M, Mouraux A. Tagging the neuronal entrainment to beat and meter. Journal of Neuroscience. 2011;31(28):10234–40.

94. Ericsson KA, Krampe RT, Tesch-Römer C. The role of deliberate practice in the acquisition of expert performance. Psychological review. 1993;100(3):363–406.

95. Anders E K. Deliberate practice and acquisition of expert performance: a general overview. Academic Emergency Eedicine. 2008;15(11):988–94.

96. Brown RM, Zatorre RJ, Penhune VB. Expert music performance: cognitive, neural, and developmental bases. Progress in Brain Research. 2015;217:57–86.

97. Aoki T, Furuya S, Kinoshita H. Finger-tapping ability in male and female pianists and nonmusician controls. Motor Control. 2005;9(1):23–39.

98. Fujii S, Kudo K, Ohtsuki T, Oda S. Tapping performance and underlying wrist muscle activity of non-drummers, drummers, and the world’s fastest drummer. Neuroscience Letters. 2009;459(2):69–73.

99. Drake C, Palmer C. Skill acquisition in music performance: relations between planning and temporal control. Cognition. 2000;74(1):1–32.

100. Repp BH. Effects of auditory feedback deprivation on expressive piano performance. Music Perception. 1999;16(4):409–38.

